# Design considerations of a wearable electronic-skin for mental health and wellness: balancing biosignals and human factors

**DOI:** 10.1101/2021.01.20.427496

**Authors:** Yasser Khan, Matthew L. Mauriello, Parsa Nowruzi, Akshara Motani, Grace Hon, Nicholas Vitale, Jinxing Li, Jayoung Kim, Amir Foudeh, Dalton Duvio, Erika Shols, Megan Chesnut, James Landay, Jan Liphardt, Leanne Williams, Keith D. Sudheimer, Boris Murmann, Zhenan Bao, Pablo E. Paredes

**Author notes:** Y.K. and M.L.M contributed equally to this work.

## Abstract

Chronic stress has been associated with a variety of pathophysiological risks including developing mental illness. Conversely, appropriate stress management, can be used to foster mental wellness proactively. Yet, there is no existing method that accurately and objectively monitors stress. With recent advances in electronic-skin (e-skin) and wearable technologies, it is possible to design devices that continuously measure physiological parameters linked to chronic stress and other mental health and wellness conditions. However, the design approach should be different from conventional wearables due to considerations like signal-to-noise ratio and the risk of stigmatization. Here, we present a multi-part study that combines user-centered design with engineering-centered data collection to inform future design efforts. To assess human factors, we conducted an *n*=24 participant design probe study that examined perceptions of an e-skin for mental health and wellness as well as preferred wear locations. We complement this with an *n*=10 and *n*=16 participant data collection study to measure physiological signals at several potential wear locations. By balancing human factors and biosignals, we conclude that the upper arm and forearm are optimal wear locations.

---

Daily stress is defined as the routine challenges of day-to-day living. These challenges can either be predictable (*e*.*g*., daily commutes) or unpredictable (e.g. a sudden deadline) and occur in 40% of all days.^[1]^ Daily stress has been shown to cause psychological distress and exacerbate symptoms of existing physical health conditions.^[2]^ Repeated trig gering of daily stress can also lead to chronic stress, which has been associated with a variety of pathophysiological risks—conditions that impair quality of life, shorten life expectancy, and can include developing mental illness.^[2, 3]^ Six hundred million people are devastated by depression and anxiety, and it is the cause for the loss of trillions of dollars each year from our global economy.^[4]^ Mental illness is now the number one silent killer of adults, and the number one cause of disability worldwide.^[5]^ According to the World Health Organization, one person dies by suicide every 40 seconds.^[6]^ Despite this crisis, available resources and access to care scarcely begin to meet the need. Complicating matters further, we have no objective tests or scalable technologies for detecting chronic stress, the type of mental illness a person is at risk for, what stage of illness they are in, nor do we know how to best intervene.

Toward addressing these needs, one promising area of research focuses on continuous sensing of physiological data using wearable sensors and devices. Wearable devices can provide unobtrusive and non-invasive monitoring of health markers making them ideal platforms for mental health and wellness monitoring. A growing body of literature indicates physiological parameters such as heart rate variability (HRV),^[7–9]^ and skin conductance (SC),^[10–12]^ and biochemical signals, such as cortisol^[13–16]^ are linked to stress, anxiety, and depression. HRV and SC are normally collected with large desktop signal acquisition units, while cortisol levels in bodily fluids are measured using enzyme-linked immunosorbent assay (ELISA)^[17]^ and liquid chromatography/mass spectrometry (LC/MS) in lab settings. With exciting advancements in electronic-skin (e-skin) and wearable technology, it is now possible to design wearables that can easily measure HRV,^[18, 19]^ SC,^[20, 21]^ and potentially corti sol.^[22, 23]^ Such a wearable can potentially enable a better understanding of how these parameters are linked to chronic stress, anxiety, and/or depression thus allowing users and their health providers to detect the onset of related mental health issues for earlier treatment and intervention. Currently, wearables are widely used for lifestyle (*e*.*g*., fitness) and medical monitoring.^[24–26]^ In these wearables, the biosignals dictate design choices while form factor is often a secondary concern. However, in the case of wearables for mental health and wellness that may be used widely by people and patients, both biosignals and human factors are important to consider to improve long term adherence when used for proactive, preventative, and treatment purposes.

Here, we present an approach that combines user-centered design with engineering-centered biosignal measurement to identify optimal wear locations for designing mental health and wellness wearables that take into account both biosignals and human factors. In our multi-part user centered design study, we first examined usability factors such as comfort, placement, and ease-of-use through a design probe study (*n*=24) that utilized a low-fidelity e-skin wearable prototype. This first component of the study investigated user perceptions and preferences of e-skin wearables for mental health and wellness applications, identified several factors that may contribute to acceptance and adherence, and explored how these perceptions and preferences might change after a short wear session using a follow-up survey. We then performed a complementary on-body data collection study to measure HRV (*n*=10), SC (*n*=10), and cortisol levels (*n*=16) at several of these potential body locations. While the wrist and the forehead are rich for sensing, users tend to prefer more discreet wear locations for privacy, such as the upper arm and torso. Thus, we used a weighting mechanism to merge both human factors and biosignals. This weighting yielded the upper arm as the optimal wear location, followed by the forearm, for e-skin mental health and wellness wearables.

Our results also suggest that wearable technologies could be adopted by end-users for not only treatment but also proactive mental wellness applications like the daily monitoring of stress. Interestingly, participants proposed such adoption could have the added benefit of normalizing conversations around mental health and wellness. However, participants remained concerned about such technologies marking them as part of a stigmatized group. As a result, factors such as comfort, size, and concealability were viewed as critical to adoption and factored into their choice in where to wear our low-fidelity wearable prototype during their short exposure.

## Design criteria of a wearable for mental health and wellness

To increase adoption, the following desired properties, as shown in Fig. 1a, should be considered during the design process of the wearable. If a sensor is imperceptible, *skin-like*, and seamless to use then there is a greater chance of adoption. Additionally, the device should not hinder the movement or comfort of the user. Privacy is another key factor that should be considered during the design process. The wearable should be *private* and concealable under everyday clothing. Since we want to get an overall snapshot of the wearer’s state of mind, the device should be *multi-sensory*. HRV, SC, and cortisol sensing capabilities are highly desirable. Furthermore, a *personalized* approach should be taken to customize the design, software, and hardware to address the needs of different individuals. Finally, to ensure personal hygiene, data quality, and convenience, the wearable should be low-cost and *disposable*.

**Figure 1.**
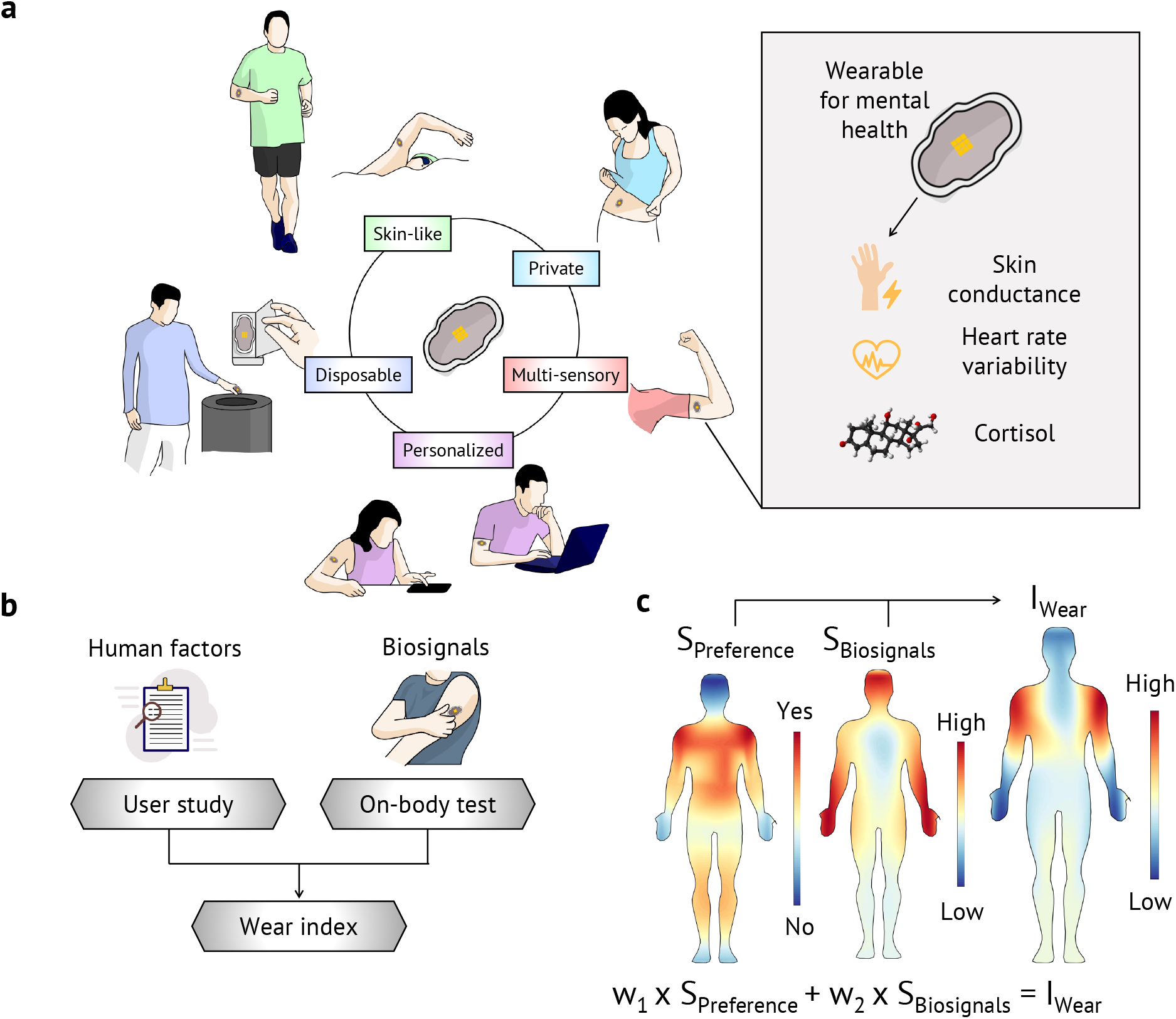
Design criteria for a mental health and wellness wearable. (**a**) Desired properties of the device: (i) The sensor should be *skin-like* and imperceptible to the user. (ii) Due to the sensitive nature of the device and data, the sensor should be *private*. (iii) The device needs to be *multi-sensory*, and collect the necessary physiological biosignals, namely, skin conductance, heart rate variability, and cortisol levels. (iv) To take a precision psychiatry approach, the device should be *personalized*, tailored to each individual, and use case. (v) To ensure reliable sensor operation and ease-of-use, the wearable should be low-cost and *disposable*. (**b**) Overview of the design approach used in this study. We collected user feedback and preference data from a *n*=24 participant study. We also performed on-body sensing to assess the quality of the biosignals at the preferred body locations. Then we weighted both human factors and biosignal qualities to create a wear index for different wear locations on the body. (**c**) Visual overview of estimating the optimal wear location. User preference data (*S*_*Preference*_) and biosignal data (*S*_*Biosignals*_) are used to find the optimal wear locations (*I*_*wear*_) on the body.

Existing wearables in commercial and academic domains are designed mostly by focusing on biosignal quality. For example, the electrocardiography (ECG) signal is the most important factor for an ECG patch. While biosignals are very important, it is necessary to include the human factors in the design process to address privacy concerns. In this work, we used both human factors and biosignals for our wearable (Fig. 1b). We studied user perception and preference (*S*_*Preference*_) on the wearability of such a sensor through a design probe study and collected biosignals (*S*_*Biosignals*_) through a lab-based data collection study. We weighted both *S*_*Preference*_ and *S*_*Biosignals*_ using different weights to reveal optimal wear locations on the body using a wear index created using a weighting mechanism: (*I*_*Wear*_ = *w*_1_×*S*_*Preference*_+*w*2× *S*_*Biosignals*_). Human factors are expressed in *S*_*Preference*_, while *S*_*Biosignals*_ expresses the contribution from the biosignals. Fig. 1c visually shows how *S*_*Preference*_ and *S*_*Biosignals*_ are utilized to find the optimal wear location.

## Human factor considerations in mental health and wellness wearable design

In our *n*=24 participant design probe study, we investigated prior experience with wearable devices as well as perceptions and preferences of a future e-skin mental health and wellness wearable (Supplementary Note 1 and Supplementary Figs. 1-7). When asked about their prior experience with wearable technology, we found that a majority of participants (87%, 21/24) strongly associated wearables with wrist-worn technology for fitness tracking, in particular, with smartwatches. A third (33%, 8/24) defined wearables as devices that monitor an aspect of the user’s health. Nearly half mentioned medical devices as examples of wearables including heart monitors, nicotine patches, and hearing aid devices. While some (17%, 4/24) had previously worn wearables for fitness tracking or medical reasons, only a small fraction (8%, 2/24) reported that they used a wearable at the time of the interview. A majority (75%, 18/24) noted that they did not need a wearable device, suggesting that they did not see a utility in them that was not covered by other common devices like their smartphones. Participants also reported high cost and lack of comfort as barriers to ownership. Of the few who were using a wearable device, most cited utility and comfort as their top criteria in selecting their wearables.

While a relatively novel use case, most participants (58%, 14/24) expressed general interest in wearables for mental health and wellness monitoring. A majority (79%, 19/24) said they would be more likely to use an e-skin wearable to measure their stress levels if it was recommended by their doctor. Those who were opposed (21%, 5/24) said medical advice would not impact their decision.

We used paper body maps (Supplementary Fig 3) and a low-fidelity version of our wearable device in the design probe study (Fig. 2a) to assess where participants might wear the e-skin. This low-fidelity device was similar to the wearable used to collect biosignals (Fig 2b) in terms of size, shape, and weight as well as the planned method of attach196 ment (*i*.*e*., using medical grade tape). In terms of where future users might wear such a device, participants showed a strong preference for the upper arms and upper torso (*i*.*e*., chest and back) followed by the stomach, waist, and thighs (Figs. 2c,d). Participants reported that concealability and comfort were the top decision factors. Thus, we note that all these body locations are usually covered by everyday clothes (*e*.*g*., t-shirt, shorts). On the other hand, visible locations such as the head and extremities (*i*.*e*., hands, wrists, and feet) were undesirable. Similarly, they disliked locations where the placement of the wearable would interfere with the body’s natural movement (*e*.*g*., elbows, knees). A condensed version of the body map results is shown in Figs. 2c,d. The complete set of results are discussed in Supplementary Fig. 5.

**Figure 2.**
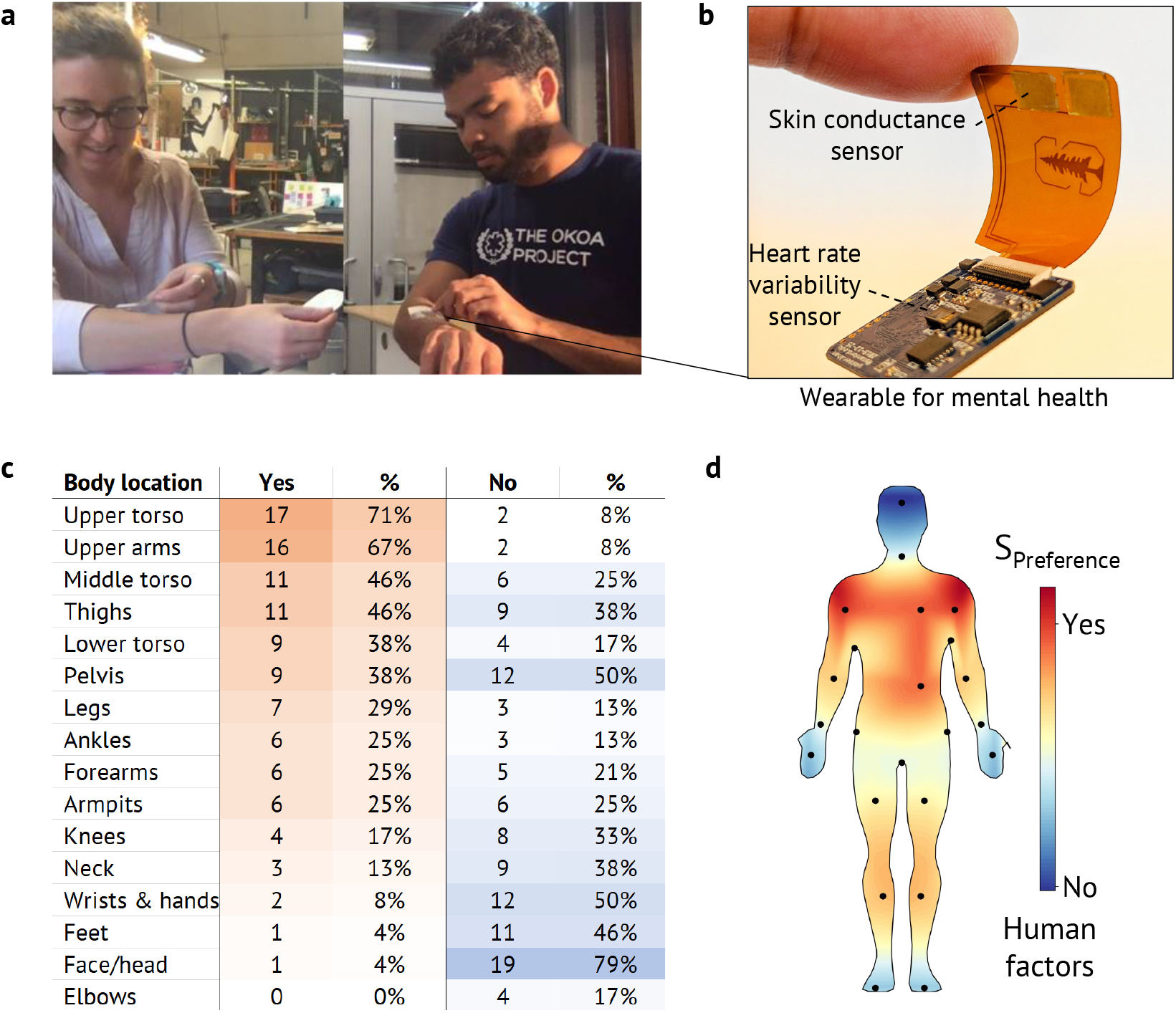
Summary of the user study on wearability, wear locations, and desired properties of a mental health and wellness wearable. **(a)** Photographs of participants interacting with low-fidelity devices with the same form factor of the developed wearable. **(b)** Sensor utilized for collecting skin conductance (SC) and heart rate variability (HRV) data. The wearable uses an optoelectronic sensor to collect HRV derived from photoplethysmography (PPG) signals. The SC data low-energy to a compatible smartphone. (**c**) Summarized table listing different body locations, positive and negative responses from the participants when asked where they would prefer to wear (Yes) or not to wear (No) the device. The presented data is condensed from the actual survey results for better understanding and visualization. The complete dataset is provided in Supplementary Fig. 5. (d) Summary data is shown visually on the body. Red regions indicate a positive preference (Yes), and blue regions indicate a negative preference (No).

When asked about how often they would change the wearable, assuming the ideal scenario where the wearable is cheap, durable, and waterproof, the answers ranged from *daily* to *monthly* with most participants preferring weekly or *bi-weekly* changes. In rationalizing these decisions, participants balanced several factors such as personal hygiene, signal continuity, convenience, and cost (Supplementary Fig. 7).

For a wearable to be socially acceptable, more than half (58%, 14/24) said its appearance is also an important factor. Participants emphasized that the ideal wearable should be fashionable (corroborating^[27]^) but also inconspicuous; it must seamlessly blend in with the rest of the wearer’s attire to avoid unwanted attention. Finally, a third said a wearable would be more acceptable if it was part of a broader social trend normalizing the management and monitoring of mental health and wellness factors. These comments are also corroborated more generally by our pre- and postsurvey results indicating that while participants were initially somewhat concerned about judgment by others or similar negative consequences of wearing such a device, they grew more positive about these concerns after a short wear session: in the post-wear survey, interest in the e-skin wearable increased and participants showed less concern that the wearable might make others uncomfortable, cause awkwardness, or result in them being ridiculed. Paradoxically, participants became more worried about what such a device might communicate about them and their iden-tity—being marked as someone in need of mental health support. The complete set of survey results are discussed in Supplementary Fig. 6.

## Biosignal measurement considerations in a mental health and wellness wearable design

Three biosignals—SC, HRV, and sweat cortisol levels are evaluated in this work. SC measures the eccrine sweat gland activity. In response to stress stimuli, a number of eccrine sweat glands get activated, and SC quantitatively measures this activity.^[10]^ HRV measures the balance between the two autonomic nervous systems—sympathetic and parasympathetic. The sympathetic nervous system gets activated when facing threats or stressors, while the parasympathetic nervous system handles the body’s relaxed state. ^[28]^ Finally, cortisol is the body’s main stress hormone. In response to internal or external stressors, cortisol is released from the adrenal glands and puts the body into a heightened-alert state. Chronic activation of the stress-response system results in overexposure to cortisol, which can disrupt almost all the body’s processes. ^[29]^ We selected sweat cortisol levels because sweat can be non-invasively collected.

Signal strengths of SC, HRV, and cortisol vary signifi pl hysmography (PPG) signal in our wearable, depends on higher the signal coming from the arteries, the better the PPG signal quality. Therefore, locations where the arteries are near the surface of the skin, provide excellent PPG sig nal. The forehead and the underside of the wrist are usually good choices for reflection-mode PPG sensing. ^[30, 31]^ On the other hand, SC depends on the density of the eccrine sweat glands, which is highest on the fingers and the palm, and drops roughly by half on the wrist and the forearm. ^[32, 33]^ We selected 6 locations on the body for a on-body data3 collection study (Fig. 3a). These locations, namely, (1) wrist, (2) forearm, (3) upper arm, (4) forehead, (5) upper chest, and (6) stomach, were chosen because of high user preference. Although the wrist and the forehead were not preferred locations indicated in the design probe study, we chose the forehead due to the high biosignal intensities, and the wrist because most commercial wearables are wrist-worn thus providing a reasonable baseline for comparison.

**Figure 3.**
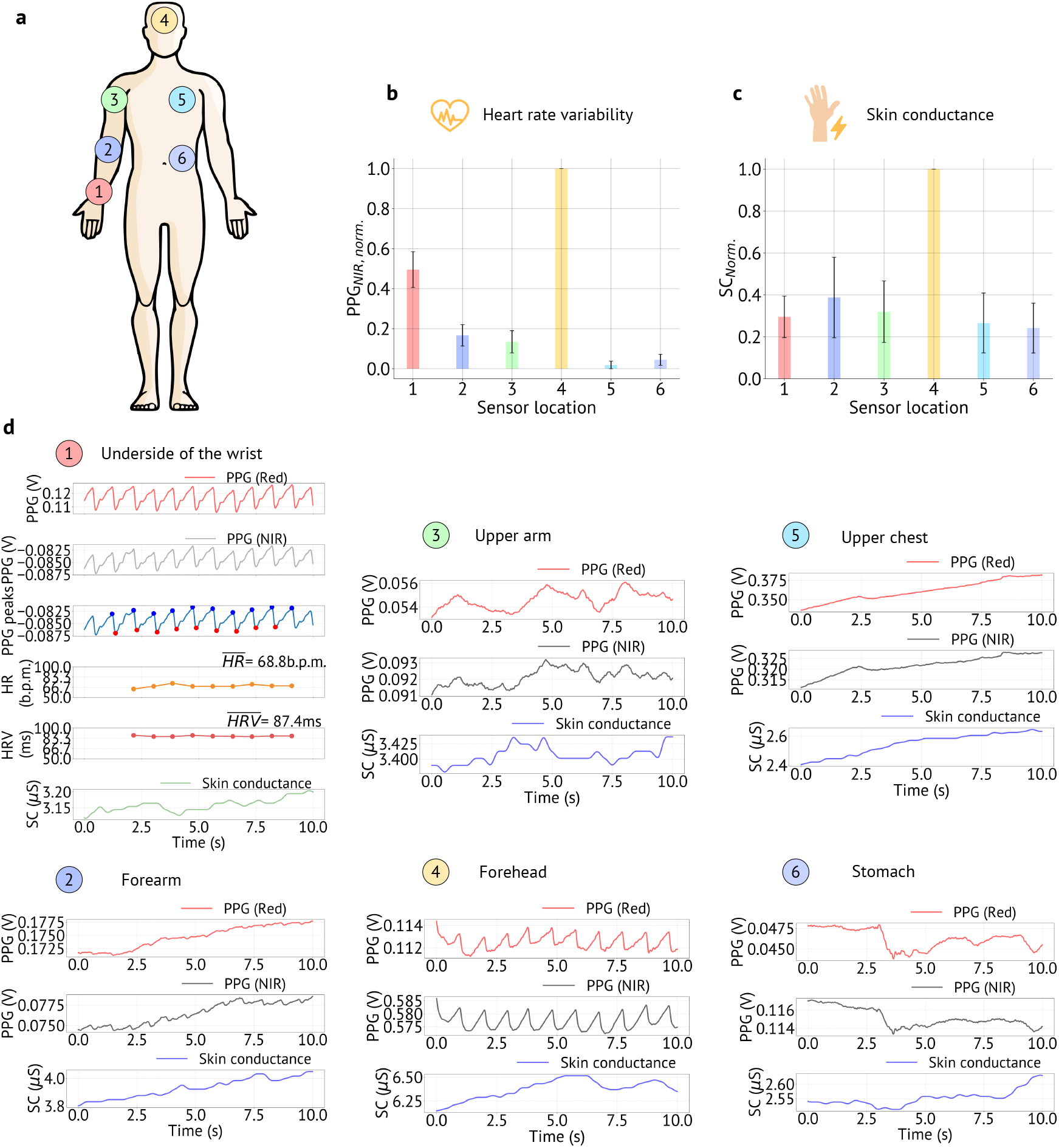
Heart rate variability (HRV) and skin conductance (SC) data distribution on the body. (**a**) Sensor placement locations − (1) wrist, (2) forearm, (3) upper arm, (4) forehead, (5) upper chest, and (6) stomach. (**b**) Photoplethymogharpy (PPG) signal magnitudes for near-infrared (NIR) light on the aforementioned 6 locations. HRV is derived from PPG, hence, PPG signal magnitudes are used in the analysis. NIR PPG signal was normalized for each participant in the *n*=10 participant study, and bar heights represent the average of the normalized value and the error bars represent the standard deviation of the normalized value. The complete dataset of *n*=10 participants is shown in Supplementary Fig. 8. The forehead shows the highest signal magnitude and gradually drops on the wrist, the forearm, and the upper arm. The signal is the lowest on the chest. (**c**) Variation of SC over the 6 highlighted locations shown in a. The SC data was normalized for each participant in the *n*=10 participant study, and bar heights represent the average of the normalized value and the error bars represent the standard deviation of the normalized value. The complete dataset of *n*=10 participants is shown in Supplementary Fig. 10. (**d**) PPG from red and NIR channels, systolic and diastolic peaks from PPG, heart rate (HR), HRV calculated from PPG signal, and SC from the 6 highlighted locations shown in a. The PPG signal is clear on the wrist, forearm, upper arm, and forehead. The PPG signal gets highly attenuated on the upper chest and stomach.

We used a custom-built wearable (Fig. 2b) to collect PPG and SC data from these 6 locations on the body. The PPG data was collected by using red and near-infrared (NIR) lights. We used the NIR PPG signal for HRV calculations. The bar chart in Fig. 3b shows the average PPG signal magnitude and variation at different places on the body. NIR PPG signal was normalized for each participant, and the average value (bar height) and the standard deviation (error bar) of the normalized data are shown in Fig. 3b. The complete dataset of *n*=10 participants is shown in Supplementary Fig. 8. The forehead provides the highest signal magnitude (100%). For NIR light, the average normalized PPG signal percentages are 49.54, 16.64,13.44, 100.00, 1.85, and 4.46 on the wrist, forearm, upper arm, forehead, upper chest, and stomach, respectively. The reproducibility of the measurement is shown in Supplementary Fig. 9, where 5 consecutive PPG measurements were collected from one participant while donning and doffing the sensor for each measurement. The upper chest showed the lowest signal magnitude and was susceptible to motion artifacts during breathing. A similar study was performed for measuring SC. We observed SC with average normalized percentages of 29.53, 38.77, 31.97, 100.00, 26.60, and 24.16 on the wrist, forearm, upper arm, forehead, upper chest, and stomach, respectively. Similar to the PPG signal, the data was collected from 10 healthy volunteers. The SC data was 307 normalized for each participant, and the average value (bar height) and the standard deviation (error bar) of the normalized data are shown in Fig. 3c. The complete dataset of *n*=10 participants is shown in Supplemntary Fig. 10. We is presented in Supplementary Fig. 11. performed a reproducibility study of the SC sensor, which is presented in Supplementary Fig. 11.

In HRV calculations, we used the root mean square successive difference (RMSSD) of the PPG signal. Five consecutive peaks were used to create a measurement window, which was moved to form a moving window for HRV calculations. Fig. 3d(1) shows the raw red and NIR PPG signals, PPG signal peaks, calculated heart rate (HR), HRV, and SC on the wrist of a volunteer. Figs. 3d(2)-(6) show the red and NIR PPG signals and SC from the forearm, upper arm, forehead, upper chest, and stomach, respectively. The PPG signal is pristine on the wrist and the forehead, but gets attenuated on the forearm and the upper arm. To calculate HRV, it is imperative that the PPG signal quality is good enough for a peak detection algorithm. Figs. 3d(1)-(4) show that the NIR PPG signals on the wrist, forearm, upper arm, and forehead are adequate for the peak detection algorithm. However, on the upper chest and the stomach, the signals barely show PPG peaks, making them unusable for HRV calculations. Both on the chest and the stomach, the PPG signals become modulated with respiration. Representative data where respiration severely affects the PPG signal is shown in Supplementary Fig. 12.

Cortisol, the third physiological parameter used in this study, was measured from sweat samples. The samples were collected from 16 volunteers at (1) forehead, (2) right arm (cubital fossa), (3) left arm (cubital fossa), (4) back of the 338 right knee (popliteal fossa), and (5) back of the left knee (popliteal fossa) (Fig. 4a). Sweat cortisol concentrations were normalized for each participant, and the average value (bar height) and the standard deviation (error bar) of the normalized data are shown in Fig. 4b. We observed average normalized cortisol percentages of 81.92, 73.19, 77.70, 344 60.96, and 59.02 on the aforementioned five locations, respectively. The complete dataset of n=16 participants is shown in Supplementary Fig. 13.

**Figure 4.**
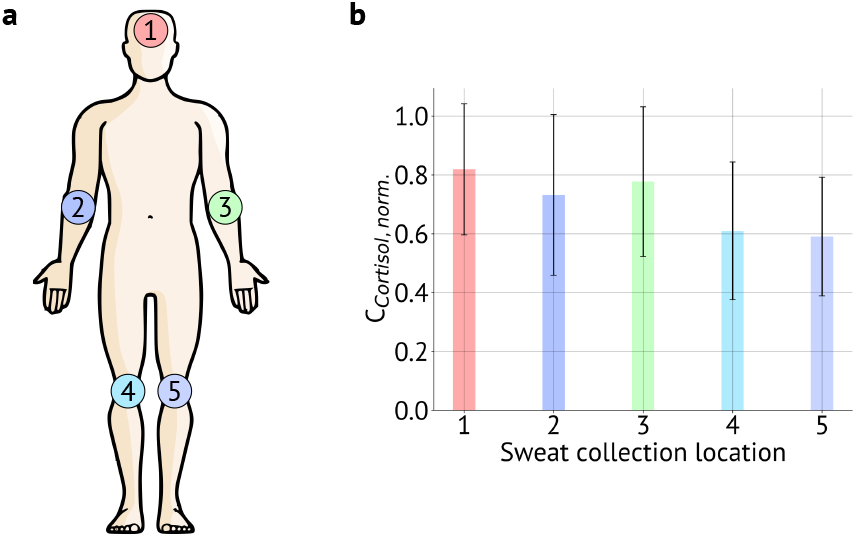
Sweat cortisol distribution on the body. **(a)** Sweat collection locations − (1) forehead, (2) right arm (cubital fossa), (3) left arm (cubital fossa), (4) back of the right knee (popliteal fossa), and (5) back of the left knee (popliteal fossa). (**b**) Sweat cortisol concentrations on the aforementioned 5 locations. Sweat cortisol concentrations were normalized for each participant in the *n*=16 participant study, and bar heights represent the average of the normalized value and the error bars represent the standard deviation of the normalized value. The complete dataset of *n*=16 participants is shown in Supplementary Fig. 13.

## Optimal placement locations for mental health and wellness wearables

To increase adoption and social acceptability, it is essential to reconcile both human factors and biosignals. In our analysis, the user preference data was collected from the design probe study, and the biosignals data was collected from the on-body sensing. For better visualization, we created body contour maps from the collected data. SC, HRV, and cortisol contour maps are shown in Figs. 5a-c. Here, the red regions signify higher signal quality, and the blue regions signify lower signal quality. The black dots represent data collection locations. All three were combined to create the biosignal body contour map using equal weights, *S*_*Biosignals*_ = *w*_1_ × *S*_*SC*_ + *w*_2_ × *S*_*HRV*_ + *w*_3_ × *S*_*Cortisol*_ where, *w*_1_ = *w*_2_ = *w*_3_ = 0.33 (Fig. 5d). The user preference body contour map was generated from the design probe study (Fig. 5e). Here, the red regions imply higher user preference, and the blue regions imply lower user preference. Finally, both human factors and biosignals were balanced to find the optimal wear location using the wear index, *I*_*Wear*_ = *w*_1_ × *S*_*Preference*_ + *w*_2_ × *S*_*Biosignals*_, as shown in Figs. 5f-h. The impact of *S*_*SC*_ and *S*_*HRV*_ on *I*_*Wear*_ is discussed in Supplementary Fig. 14. We used various weight combinations to examine the evolution of the wear location based on *S*_*Preference*_ and *S*_*Biosignals*_. When the preference data is weighted highly at *S*_*Preference*_ = 75% and *S*_*Biosignals*_ = 25%, the *I*_*Wear*_ is high at locations that are generally hidden under clothing (Fig. 5f). In the opposite case, when the biosignals are weighted heavily at SBiosignals = 75% and *S*_*Preference*_ = 25%, the *I*_*Wear*_ is high at the extremities of the body such as the forehead or the 378 wrist (Fig. 5h). When both user preference and biosignals are balanced at *S*_*Preference*_ = 50% and *S*_*Biosignals*_ = 50%, a compromise is reached, and *I*_*Wear*_ is high on the upper arm and the forearm. Hence, the upper arm or the forearm is the optimal sensing location for our e-skin wearable, where the biosignals are of adequate strength and the location provides privacy to the users.

**Figure 5.**
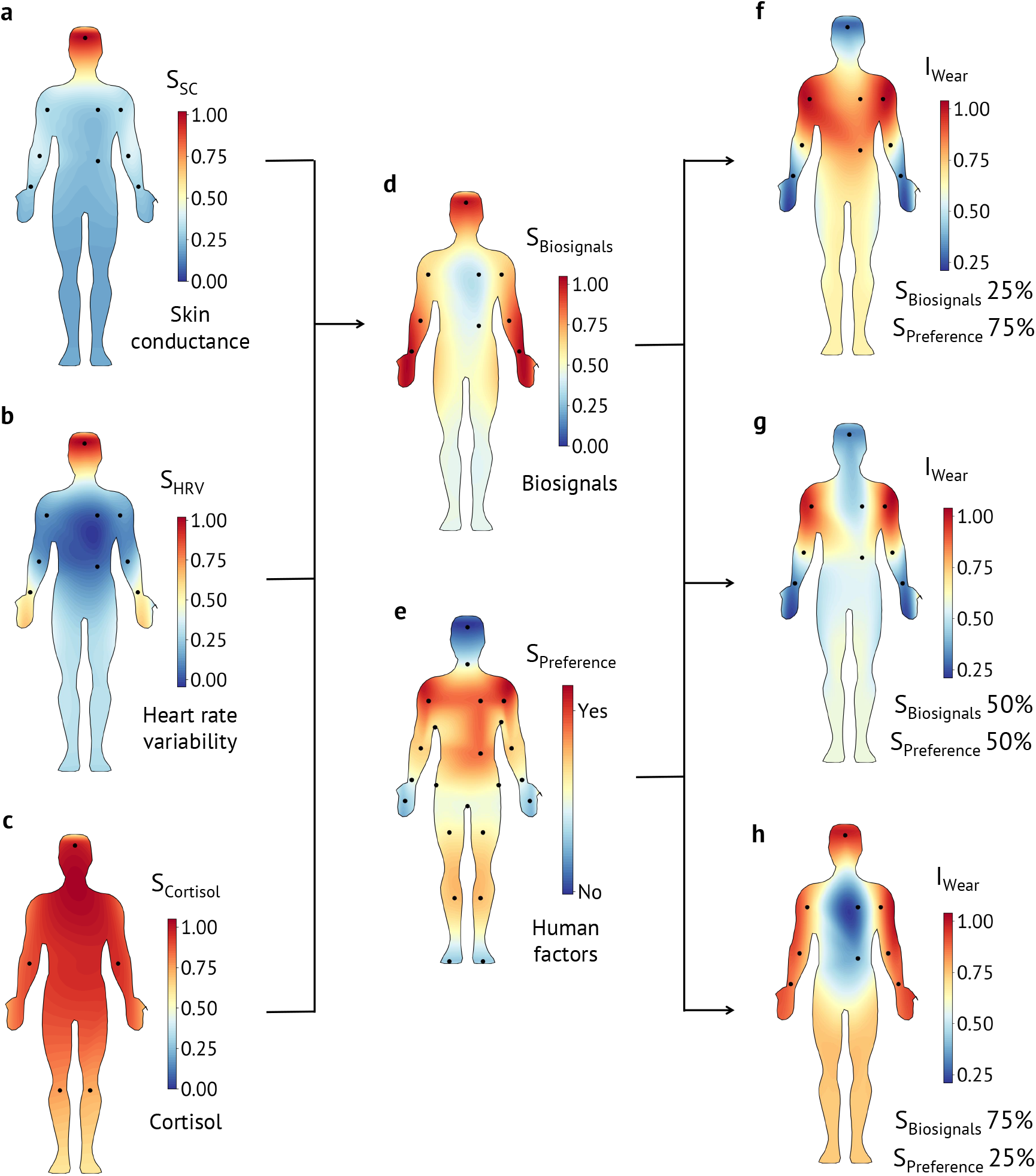
Optimal wear locations for a mental health and wellness sensor. **(a-c)** Distribution of skin conductance (SC), heart rate variability (HRV), and cortisol on the body. Red regions designate higher signal magnitude, and blue regions designate lower signal magnitude. **(d)** Distribution of combined biosignals (*S*_*Biosignals*_) on the body. SC, HRV, and cortisol signal magnitudes are equally weighted to generate the contour map. **(e)** User preference data (*S*_*Preference*_) is shown using a contour map on the body. Here, red regions show a positive preference, and blue regions show a negative preference. (**f-h**) Optimal wear locations are shown using the wear index (*I*_*Wear*_). Red regions show a high *I*_*Wear*_ and blue regions show a low *I*_*Wear*_. Three different weight combinations are used to generate contour maps. In f, user preference is weighted heavily at *S*_*Preference*_ = 75% and *S*_*Biosignals*_ = 25%. It is evident that using user preference, the wear locations are mostly hidden under the clothing on the upper body. In g, both *S*_*Preference*_ and *S*_*Biosignals*_ are weighted at 50%, which yields forearms and upper arms as the optimal wear locations. In h, the biosignals are weighted heavily at *S*_*Biosignals*_ = 75% and *S*_*Preference*_ = 25%. In this case, the optimal wear locations move to the extremities of the body where the biosignal strengths are strong.

## Conclusions

Our work corroborates aspects of prior work around factors that influence wearable design while highlighting concerns more specific to mental health and wellness applications. 388 For example, Zeagler et al. developed various body contour maps that can be used to inform wearable design noting items like motion impedance (similar to our work) as a concern or that certain areas of the body are optimal for PPG sensing.[34] However, these factors were viewed individually. Our work unifies biosignals with human factors to build a context-aware body contour map in addition to contributing body contour maps for additional sensing (*i*.*e*., SC and cortisol) and location preferences. As our context is mental health and wellness, privacy and discreetness are prioritized due to concerns around social stigmatization. ^[35, 36]^ We find 399 that these concerns may be a significant barrier to the acceptability of mental health and wellness wearables. In our study, half of the participants considered perceived judgment by others to be a downside of using one. A third were worried the wearable would distract from their daily conversations or prompt questions by others.

These social considerations are reflected in participants’ preferred wear locations and must be considered during the design process. Whereas most common health and fitness wearables are worn on the wrists, we observe that participants particularly care about discreetness of the wear location when it comes to mental health and wellness wearables. For instance, exposed body locations such as the face, hands, and wrists were among the wear locations most disliked by participants because they were perceived as distracting, uncomfortable, and public. However, when it comes to building wearables, designers are limited not only by user preferences but also by the availability of biosignals in different body locations. Since much of the wearable industry has focused on a few specific body locations (*e*.*g*., wrists), there is limited research into the availability of biosignals in other areas (*e*.*g*., upper arms, back, and chest) preferred by participants in our study. Our aim with these results is to encourage designers and researchers to develop new wearables that work on these discreet locations of the body. Thus, our findings and approach (*i*.*e*., the union of biosignals and user preferences) may serve as design guidelines for future mental health and wellness wearables.

While our work has focused on the complexiity of and potential barriers to adopting e-skin wearables, it is important to note some concerns about novelty effects when working with participants. While our participants were familiar with wearable devices, e-skin devices are still relatively new, and negative reactions to their use in public has been noted in other contexts (*e*.*g*., e-skins devices for interactions with other electronic devices^[37]^). Moreover, we derived our usability and experiential questionnaires from the WEAR Scale^[38, 39]^ to understand perceptions of e-skin wearables as this is important for early design work; however, future work should explore using a more robust (or the complete) acceptability scale when evaluating higher fidelity iterations. Finally, as far as we know all participants were healthy individuals and future work should involve patients.

## Methods

### Human factors study

We conducted a two-part design probe study to investigate users’ perceptions of wearable devices and emerging e-skin technologies for stress monitoring and other mental health applications (Supplementary Figs. 1-7). In part one, public kiosks were set up at three different locations: a campus café, the campus bookstore, and the local public library. From these kiosks, we recruited passersby for brief semi-structured interviews (*Median*=24 min, *standard deviation*=4.5 min). In addition to questions about their wearable device use, participants were asked to indicate on paper body contour maps (Supplementary Fig. 3) where they would and would not wear an e-skin for mental health and wellness applications while “thinking aloud” to explain their rationale. They then applied a low-fidelity version of our sensor to their preferred body location using medical grade tape and completed a short survey (derived from the WEAR scale^[38, 39]^) about their demographics, the comfort of the low-fidelity wearable prototype, and the perceived social acceptability around its use. In part two, we asked participants to go about the rest of their day while continuing to wear the low-fidelity wearable prototype and then to complete a follow-up survey similar to the prior but with additional open-text response questions about their experience; this data was then treated as a pre-post test with results presented in Supplementary Fig. 6.

### Human factors study participants

In total, we recruited 24 participants (12 male, 11 female, 1 non-binary) from the Palo Alto, California area. Participants were, on average, 35.8 years old (*Median*=28, *standard deviation*=15.4). Most (79%) had a high degree of formal education (bachelor’s and higher) and most (79%) were white or asian. Half (50%) were working full-time and over a third (38%) were students. Scores on the short Perceived Stress Scale (PSS-4)^[40, 41]^ indicate that most experienced moderate levels of stress over the last month (*Median*=6.44, *standard deviation*=3.29) (Supplementary Fig. 2). All experiments were performed in strict compliance with the guidelines of IRB and were approved by the Committee for Protection of Human Subjects at Stanford University (protocol no., IRB-45825). Informed consent was obtained from all participants.

### Human factors data and analysis

In sum, data from this study includes: survey responses, paper body contour maps, and interview transcripts. Descriptive statistics were calculated from closed-form survey results while open-response questions were thematically analyzed. Similarly, descriptive statistics were generated about regions indicated on the paper body contour maps. All interviews were recorded and professionally transcribed for computer-assisted qualitative data analysis using NVivo (v12). A researcher began the analysis by designing a preliminary codebook based on our research questions as well as concepts raised in prior literature. Random selections of 12% of the interview transcripts were independently coded by two researchers according to this primary codebook and inter-rater reliability (IRR) was measured using Cohen’s kappa (*κ*). Between rounds, the researchers met to resolve disagreements and update the codebook. An overall *κ*=0.83, considered an almost perfect agreement, was achieved after two rounds of coding. The remaining interviews were then independently coded.

### Biosignal data collection and processing

#### SC and HRV data collection study

SC and HRV data collection were performed using a custom-built wearable device. In the sensor, a pair of electrodes with hydrogel was used to collect the SC data. Using a feedback loop with a pair of operation amplifiers (op amps), we ensured that *<*10*μ*A current flows for typical SC in the range of 0-50 *μ*S. Texas Instruments TLV9102, dual 1MHz, 16-V rail-to-rail op amps were used to implement the SC read- out circuit. The output signal was sampled using a 12-bit analog-to-digital-converter (ADC) of a Nordic Semiconductor nRF52832 Bluetooth transceiver.

The HRV signal was obtained from PPG signals collected by an optical sensor. SFH 7050 from OSRAM Opto Semiconductors Inc. was interfaced with the nRF52832 Bluetooth transceiver using a serial peripheral interface (SPI). Red (660 nm) and NIR (950 nm) lights were used to collect the PPG signals at 100 Hz sampling frequency. A silicon photodiode of the SFH 7050 sensor was used to collect the reflection-mode optical signal.

#### SC and HRV data collection study participants: 10

healthy volunteers (6 male, 4 female) participated in the on-body SC and HRV data collection study. The volunteers were asked to put on the sensors on 6 different locations on the body. Then SC and HRV data were collected using the wearable and a mobile app for 2 minutes at every location. All experiments were performed in strict compliance with the guidelines of IRB and were approved by the Committee for Protection of Human Subjects at Stanford University (protocol no., IRB-41837).

### SC and HRV data analysis

SC raw data was collected from the sensor and sent over Bluetooth to a smartphone. In the case of HRV, PPG signals from red and NIR channels were collected, and the NIR signal was used in a peak detection algorithm to find the systolic peaks. HR and HRV were calculated from the systolic peaks. RMSSD of the peaks, 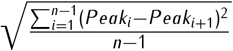 were used to calculate the HRV. Here, five consecutive systolic peaks (n=5) were used 542 to create a windowed measurement.

### Cortisol data collection study

Sweat cortisol samples were collected from volunteers during a body temperature manipulation study, which was part of a larger protocol. Volunteers sat in a portable dry infrared sauna that zipped up around the chin. Their whole body was enclosed in the sauna except their head. The sauna temperature was set to 60 °C (140 ° F). Volunteers remained in the sauna until either 45 min had elapsed, or until their core body temperature reached the maximum safety limit of 39.4 °C (103 ° F). 552 Volunteers had their core body temperature measured using an infrared tympanic membrane thermometer every 3 min that they were in the sauna to ensure that their core body temperature did not get too high. We collected sweat samples from participants as their bodies attempted to regulate their core body temperature. Sweat was collected utilizing an array of non-woven dental sponges to absorb the sweat from the skin surface. Dental sponges were affixed to the body using a transparent stretchable and waterproof medical dressing (Tegaderm, 3M). Sweat was collected from the forehead proximal to the frontal bone, the cubital fossa (inside of elbow), popliteal fossa (back of the knee). The cubital fossa and popliteal fossa dental sponges were placed bilaterally on both the left and right sides. Once volunteers exited the sauna the sweat saturated dental sponges were placed in centrifuge-compatible tubes originally designed to extract saliva from cotton swabs (Salivette system, Sarstedt,inc). The dental sponges were centrifuged at 3300 revolutions per minute (rpm) for 10 min to separate sweat from the dental sponge. Sweat samples were then frozen and stored at −80 ° C until they were thawed for analysis.

### Cortisol data analysis

The analysis of sweat samples was conducted by Dresden lab service utilizing a standard 574 ELISA with a 0.2 nmol limit of detection (LOD) and a co-efficient of variability of<7% for both the inter-assay and intra-assay measures.

### optimal placement location

In the optimal sensor placement analysis, the biosignal data for SC, HRV, and cortisol were normalized first using the equation: 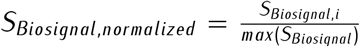. To compute the overall effects of biosignals, SC, HRV, and cortisol data were equally weighted using the equation: *S*_*Biosignals*_ = *w*1 × *S*_*SC*_ + *w*_2_ × *S*_*HRV*_ + *w*_3_ × *S*_*Cortisol*_, where, *w*_1_ = *w*_2_ = *w*_3_ = 0.33. Throughout this work, the contour maps were generated by interpolating the sensor data in 2D space. A false average color was assigned to the corners of the plots for better visualization. After that, both human factors and biosignals were used to generate the wear index, *I*_*Wear*_ = *w*_1_ × *S*_*Preference*_ + *w*_2_ × *S*_*Biosignals*_. Here, *w*_1_ and *w*_2_ were assigned the combinations of (*w*_1_ = 0.75, *w*_2_ = 0.25), (*w*_1_ = 0.50, *w*_2_ = 0.50), and (*w*_1_ = 0.25, *w*_2_ = 0.75) to investigate the effects of human factors and biosignals in determining the optimal sensor placement. All analyses were performed using custom-written Python 3.6 scripts

## Acknowledgements

This work was supported in part by the Stanford Catalyst for Collaborative Solutions and the National Science Foundation SenSE program (grant no. 2037304). We thank Min-gu Kim, Chenxin Zhu, Hongping Yan, Sahar Harati, and Eusha Abdullah Mashfi for technical discussions and helping with the manuscript. Contributions from M.L.M. were made while in transition from Stanford University to the University of Delaware.

## Author contributions

Y.K., M.L.M., Z.B., and P.E.P. designed the research. Y.K.,N.V., J.Li, J.K., A.F., D.D., and E.S. contributed to the biosignals portion of the study. M.L.M., P.N., A.M., and G.H. contributed to the human factors portion of the study. J.Landay, J.Liphardt, L.W., K.S., B.M., Z.B., and P.E.P. oversaw the project. Y.K., M.L.M, P.N, Z.B., and P.E.P. wrote the manuscript, and all authors edited the manuscript

## Additional information

### Supplementary Information

accompanies this paper at [Link]

### Reprints and permission

information is available online at

**How to cite this article**

## Supplementary information

### 1 Supplementary Note 1: Human factors study

We conducted a two-part design probe study to investigate users’ perceptions of wearable devices and emerging electronic-skin (e-skin) technologies for stress monitoring and other mental health and wellness applications (Supplementary Figs. 1-7). In part one, public kiosks were set up at three different locations: a campus café, the campus bookstore, and the local public library. From these kiosks, we recruited passersby for brief semi-structured interviews (*Median*=24 min, *standard deviation*=4.5 min). In addition to questions about their wearable device use, participants were asked to indicate on paper body diagrams (Supplementary Fig. 3) where they would and would not wear an e-skin for mental health and wellness applications while “thinking aloud” to explain their rationale. They then applied a low-fidelity version of our sensor to their preferred body location using medical grade tape and completed a short survey about their demographics, the comfort of the wearable prototype, and the perceived social acceptability around its use (derived from the WEAR Scale^[1, 2]^). In part two, we asked participants to go about the rest of their day while continuing to wear the low-fidelity prototype and then to complete a follow-up survey similar to the prior but with additional open-text response questions about their experience.

We recruited *n*=24 participants (12 male, 11 female, 1 non-binary) from the Palo Alto, California area. Participants were, on average, 35.8 years old (*Median*=28, *standard deviation*=15.4). Most (79%) had a high degree of formal education (bachelor’s and higher) and most (79%) were white or asian. Half (50%) were working full-time and over a third (38%) were students. Scores on the short Perceived Stress Scale (PSS-4)^[3, 4]^ indicate that most experienced moderate levels of stress over the last month (*Median*=6.44, *standard deviation*=3.29) (Supplementary Fig. 2). When asked about their prior experience with wearable technology, we found that a majority of participants (87%) strongly associated wearables with wrist-worn technology for fitness tracking and, in particular, smartwatches. By aggregating participant’s paper body maps (Supplementary Fig 3), we are able to generate location preferences for e-skin wearables for mental health and wellness.

**Supplementary Fig 1.**
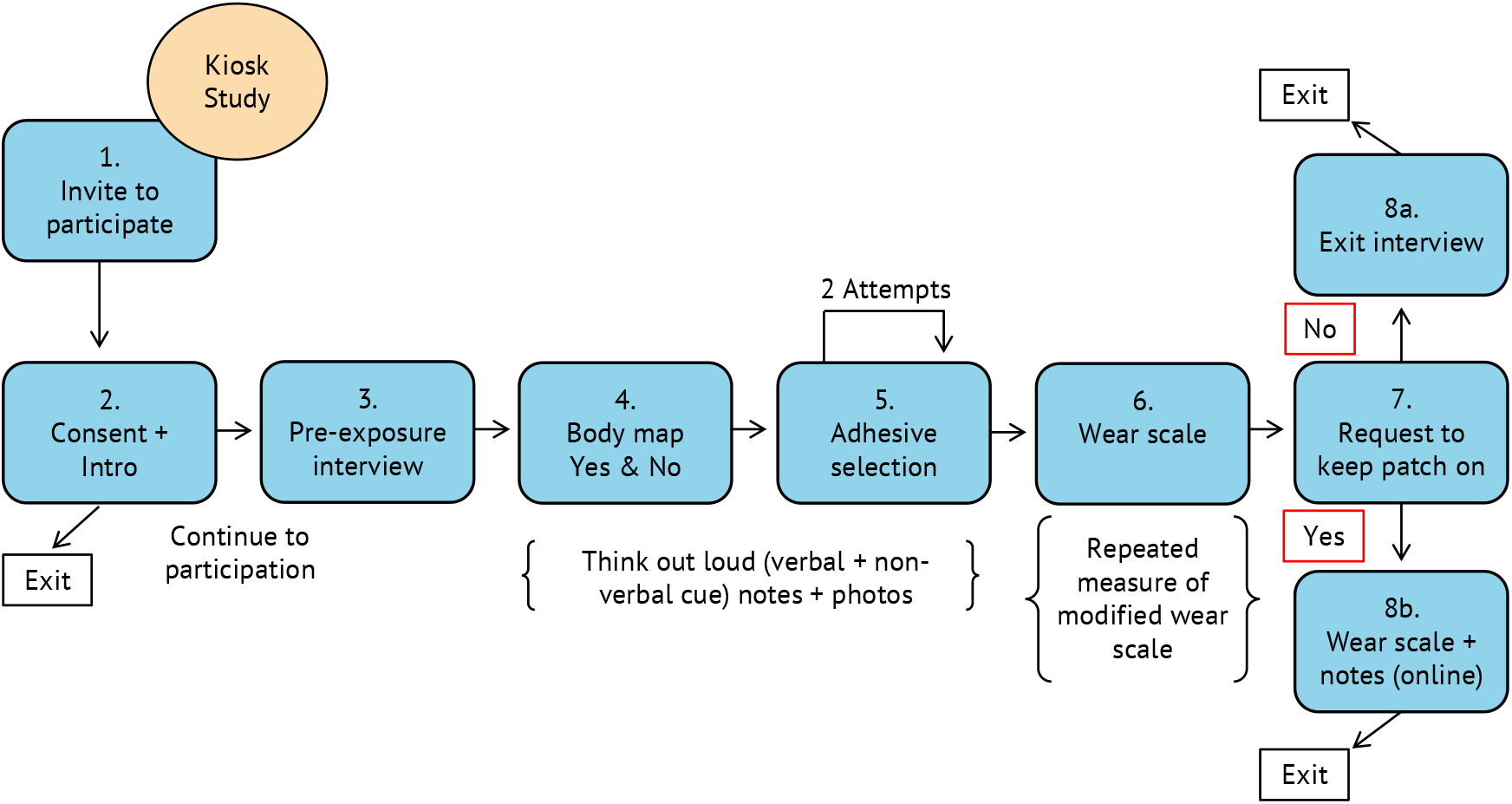
Design probe study overview. (1) Potential participants are recruited from passersby. (2) Researcher and potential participants review the consent form with the reviewer answering any questions about the process. (3) The researcher conducts a brief semi-structured interview asking about the participants prior experience with wearable devices and their thoughts on using wearables for mental health and wellness applications (*e*.*g*., monitoring daily stress). (4) Participants are then introduced to the concept of an electronic-skin (e-skin) wearable device and asked where they might be comfortable wearing such a device given the mental health and wellness context. (5) Participants are then asked to apply a low-fidelity mockup of our e-skin wearable device to their skin. (6) After adhering the mockup device, participants are asked to complete a survey that asks them about their demographics as well as questions derived from the WEAR Scale^[1, 2]^ focusing on social perceptions of e-skin wearable devices, comfort of the wearable prototype, and other factors. (7) Participants are then asked if they would mind continuing to wear the mockup device for the rest of their day. The session ends with participants either removing the mockup device (8a) or continuing to wear the device and completing another survey later that evening (8b); the follow-up survey was similar to the first with demographic questions removed and open-form experiential questions added.

**Supplementary Fig 2.**
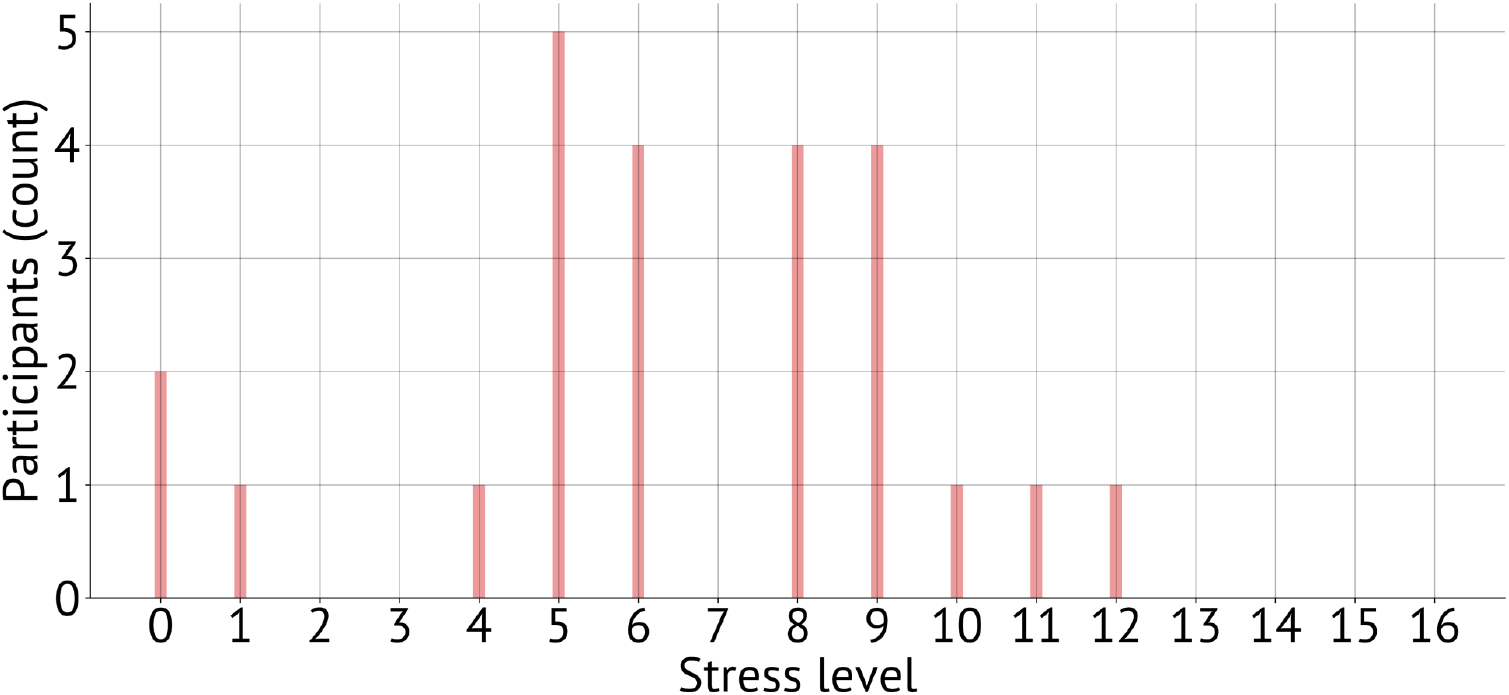
Perceived stress levels (from design probe survey) As part of the demographic questions asked on the survey during the in-person design probe sessions, participants completed the short Perceived Stress Scale (PSS-4)^[3, 4]^ to assess the level of stress they experienced in the past month. Results indicate that most participants experienced a moderate level of stress (*Median*=6.44, *standard deviation*=3.29).

**Supplementary Fig 3.**
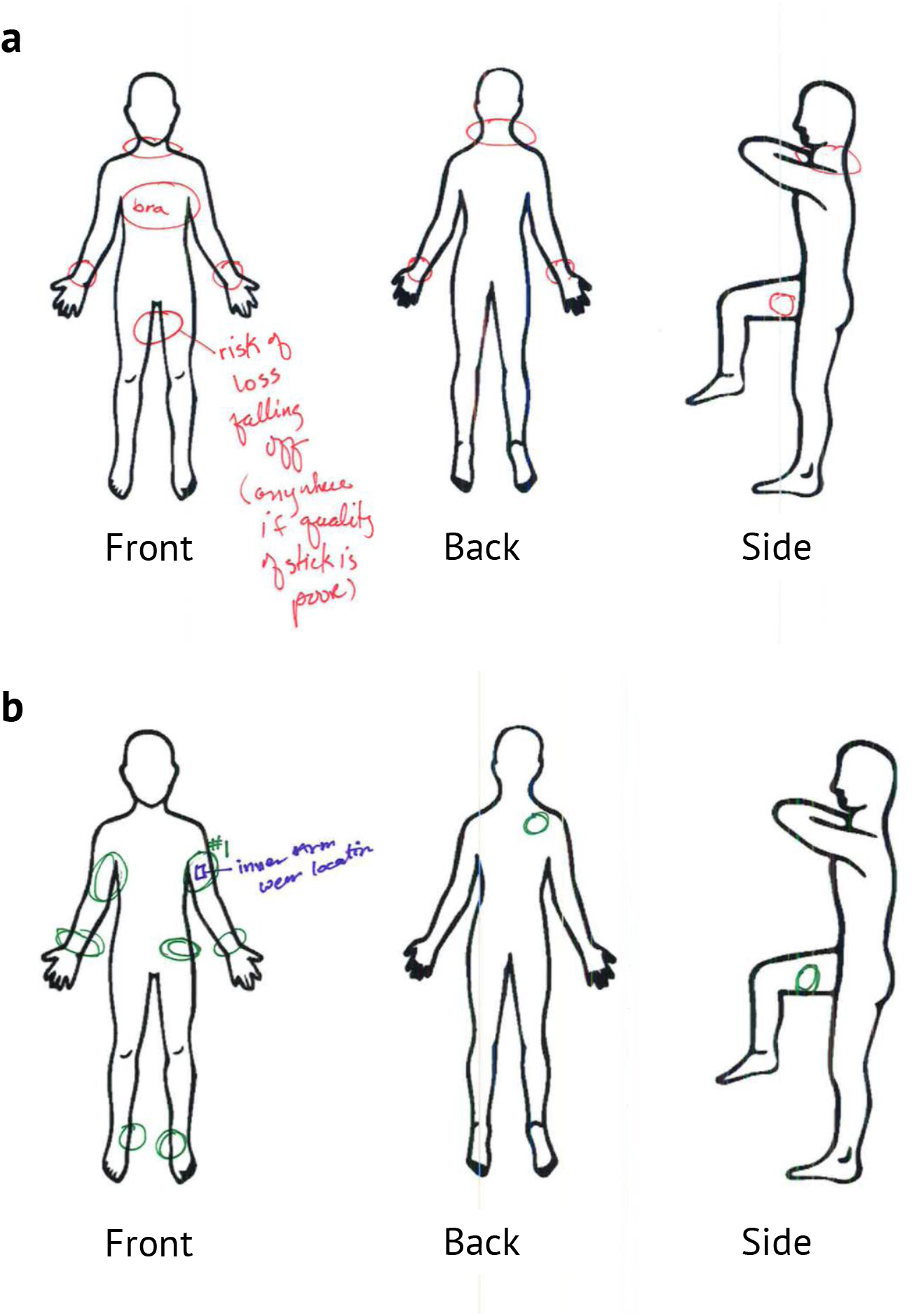
Survey body maps. To inform the future design of e-skin wearables for mental health and wellness applications, participants completed paper body maps to indicate where they would and would not be comfortable wearing an e-skin wearable. Participants were instructed to “think aloud” and annotate their body maps while the researcher collected notes. (**a**) The participant has indicated that the inside of the legs would be problematic (red) for them—worrying that the wearable may fall off as a result of natural body movement. (**b**) The participant also indicated that the upper arm was their preferred location (green) with a “#1” and that this was where they wore the low-fidelity mockup of the device (blue) during and after departing the design probe session.

**Supplementary Fig 4.**
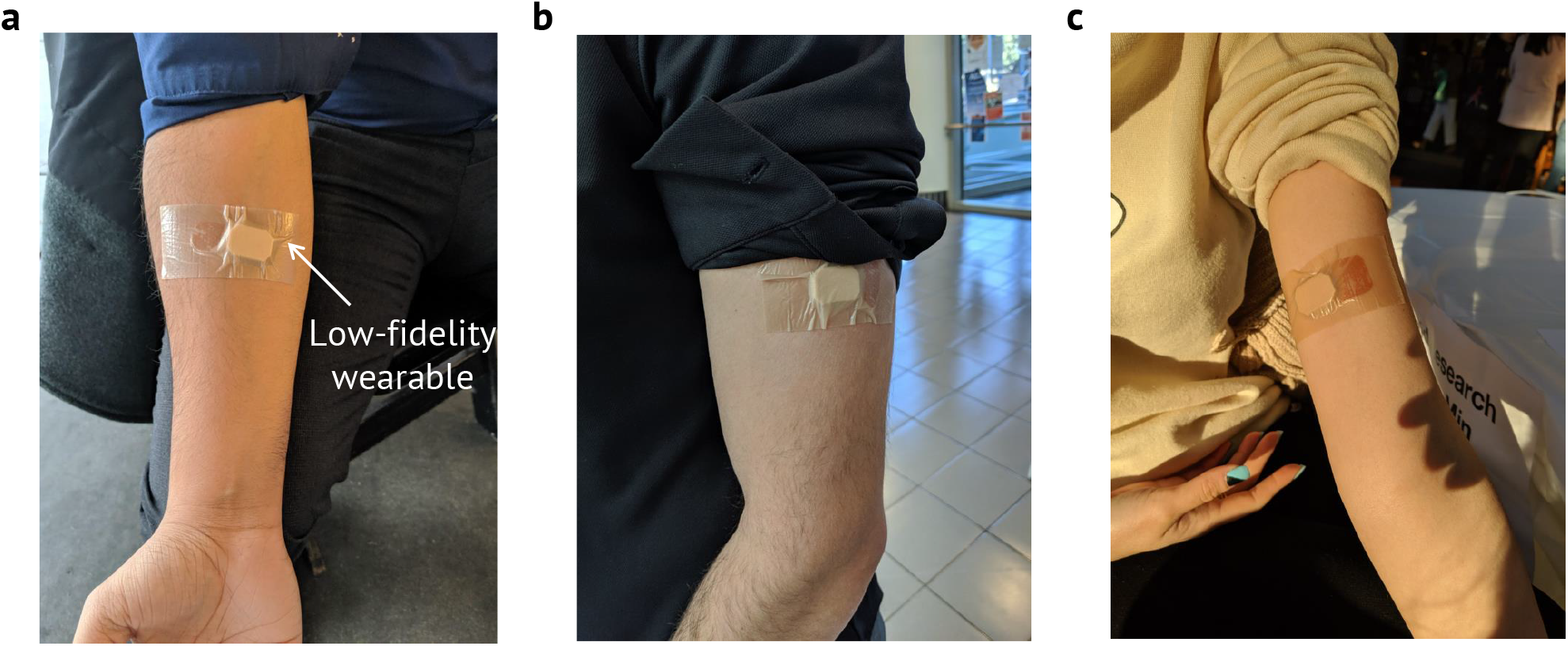
Examples of wear locations. With participant permission, the researcher at the design probe collected photos of wear locations that were often (**a**) the underside of the forearm, (**b**) the outside of the upper arm, and (**c**) the inside of the upper arm.

**Supplementary Fig 5.**
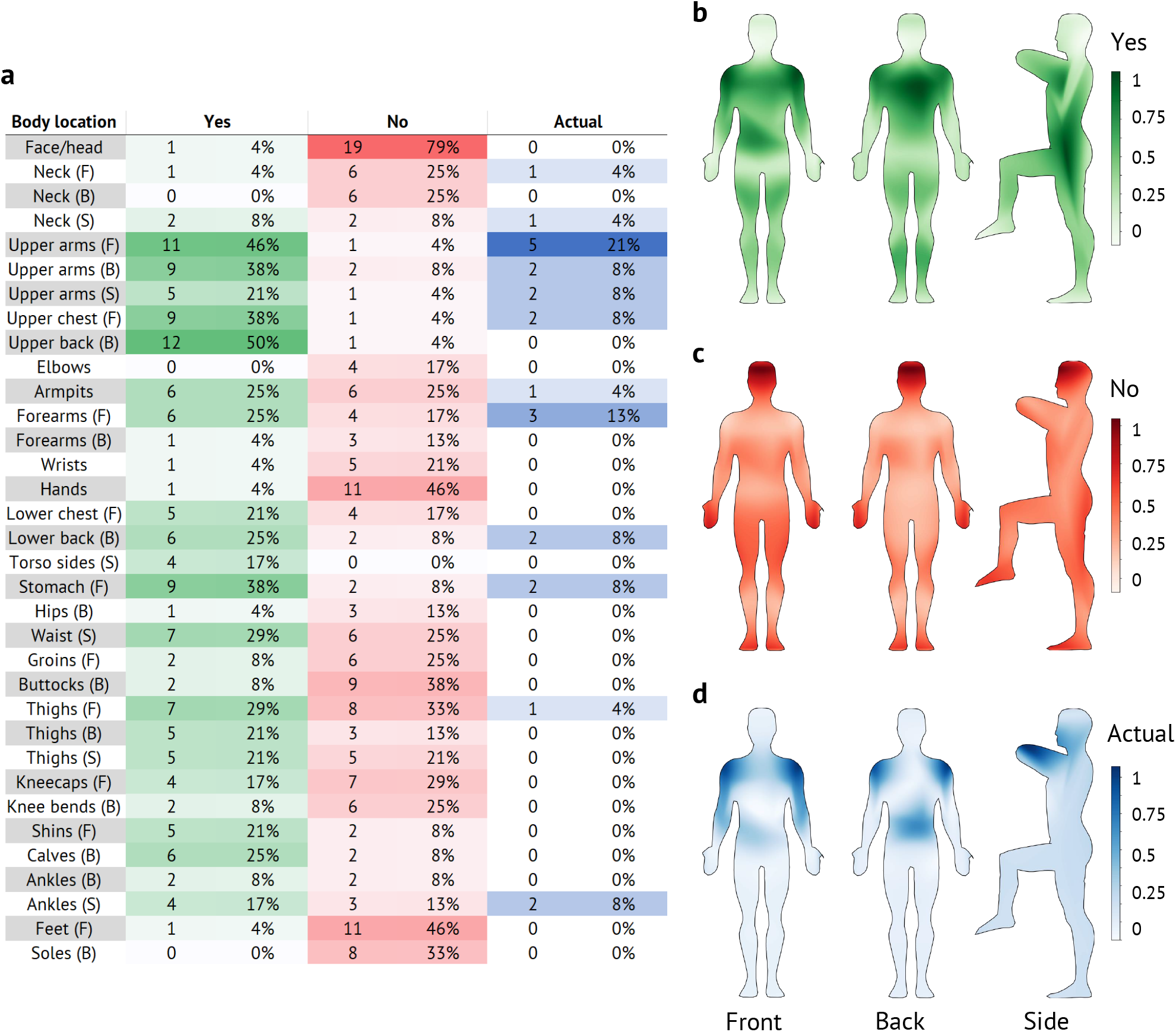
Wear locations compiled: (**a**) Body maps were divided into 34 regions and counts were aggregated based on where participants would and would not wear an e-skin device for mental health and wellness applications as well as where they actually wore the low-fidelity device during and after the design probe session. (**b-d**) This data was converted into body contour maps, which demonstrate preferences that can inform future designers of such technology.

**Supplementary Fig 6.**
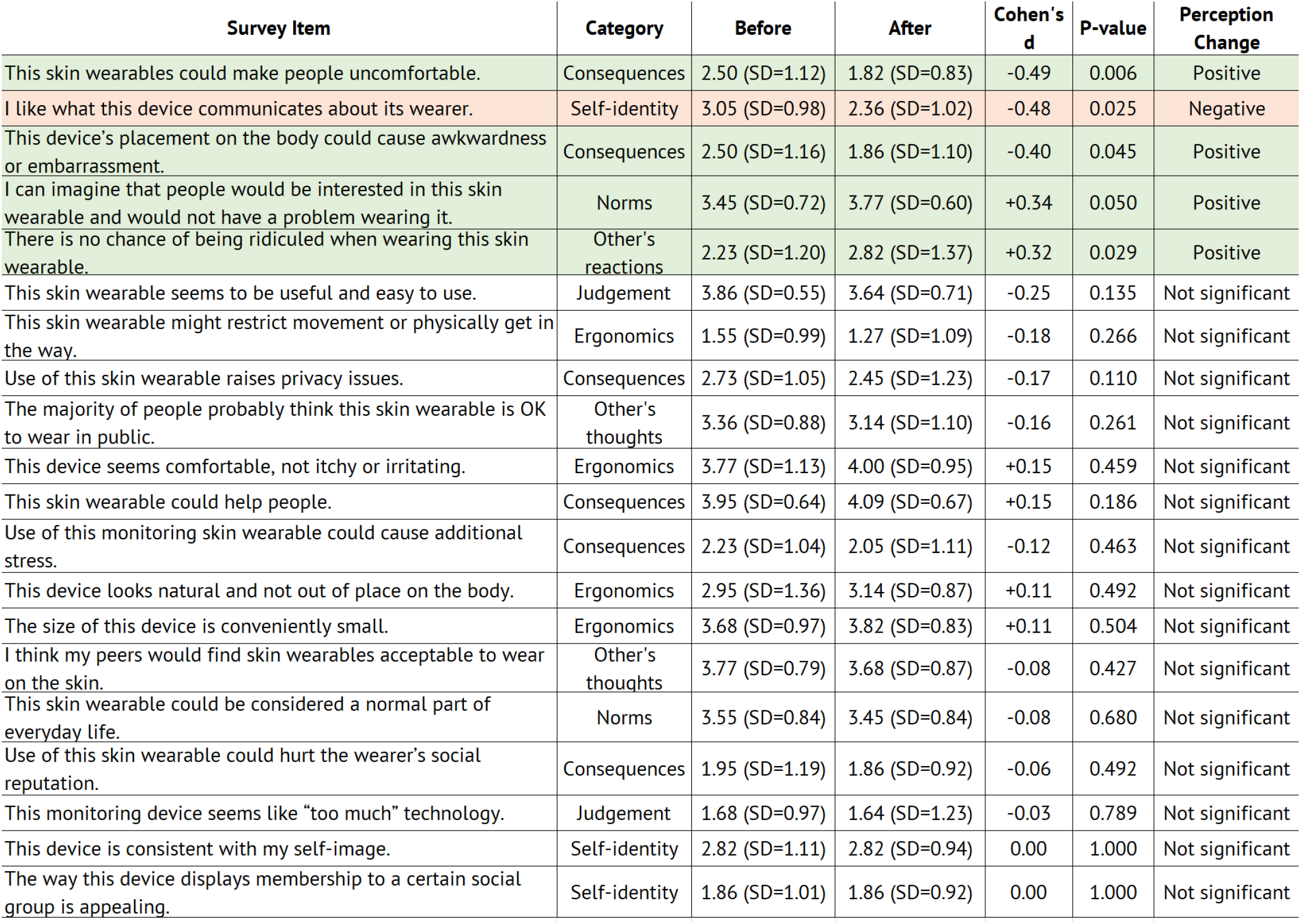
Partial WEAR Scale derived questionnaire (from design probe study survey) Participants who completed both parts of our design probe study rated several items derived from the WEAR Scale^[1, 2]^ that corresponded to social perceptions, ergonomics, and self-image. On a five point likert, higher scores tends to reveal positive attitudes while lower scores tend to indicate negative attitudes. While overall impressions of e-skin wearables were positive and consistent pre- and post-wear, we detected a significant change in 5 specific questions. We calculated the change significance using the parametric t-test after the Shapiro-Wilk test confirmed the normality of the distribution of the responses to each survey item. Additionally, Cohen’s d was calculated as a relative measure of the change size. The comparison of the pre- and post-wear survey results indicate that after a short wear of a low-fidelity device, participants showed less concern that the wearable might make others uncomfortable, cause awkwardness, or result in being ridiculed. Paradoxically, while participants believed there were less negative consequences with wearing the device, they became increasingly more worried about what such a device might communicate about them.

**Supplementary Fig 7.**
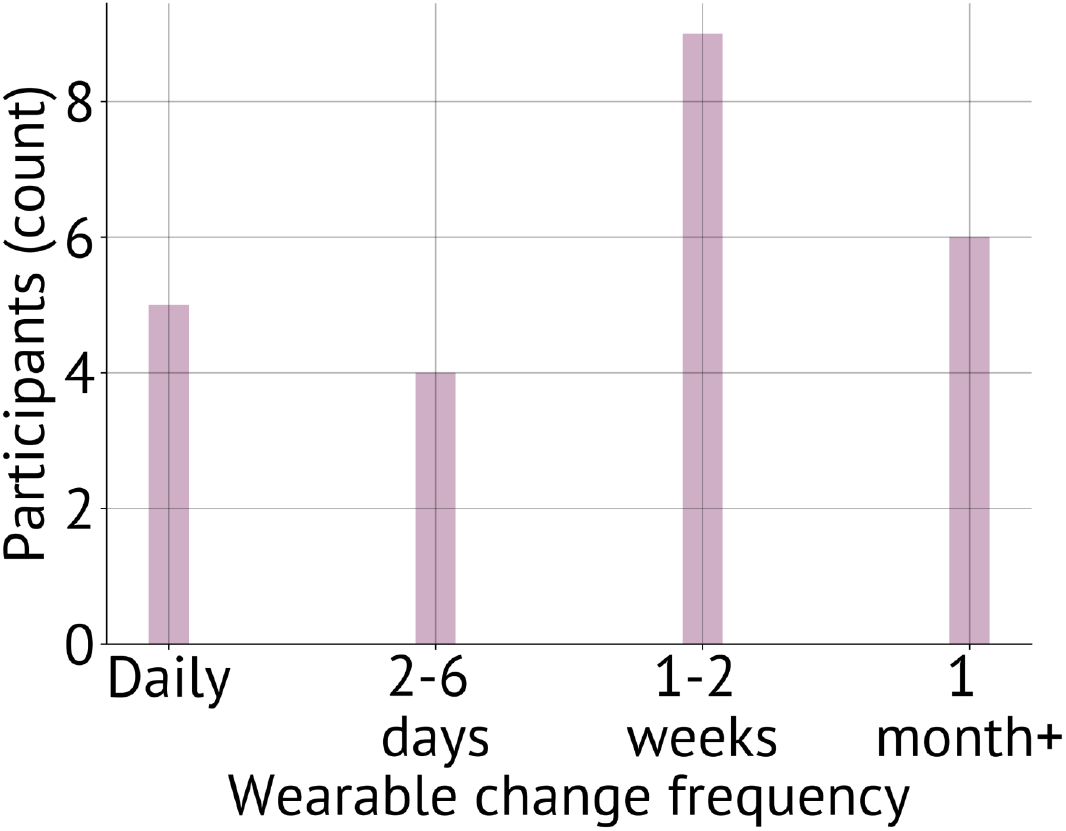
Hygiene (from design probe survey) Participants were asked how frequently they would want to change an e-skin wearable. Most would prefer to change the wearable between 1 − 2 weeks—similar to charging a smartwatch. Others would prefer to change the e-skin wearable device daily or even every few days while others would prefer to change the device every month or more (i.e., for consistent longitudinal data).

**Supplementary Fig 8.**
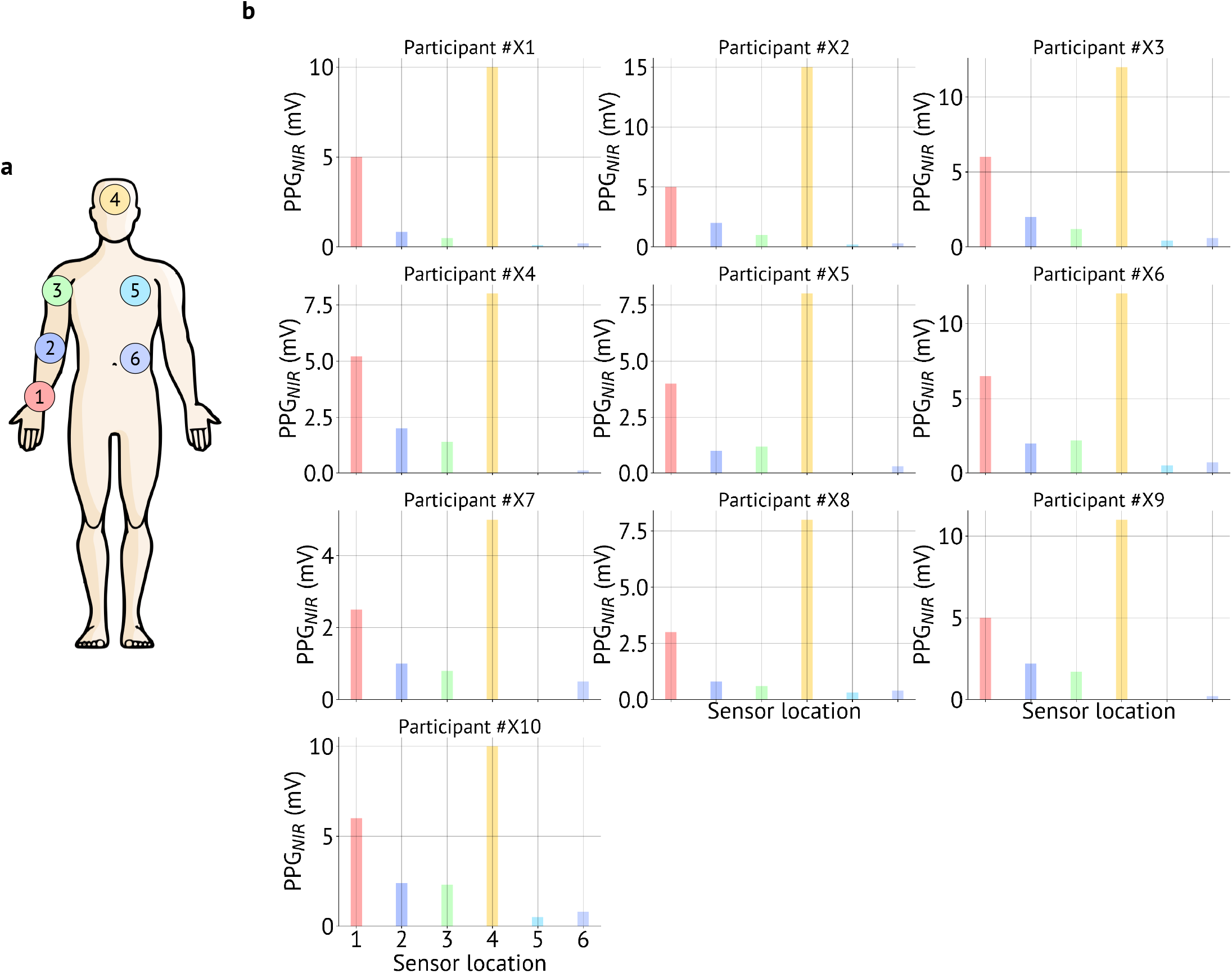
The complete NIR PPG dataset of *n*=10 participants used in the HRV analysis. (**a**) Sensor placement locations − (1) wrist, (2) forearm, (3) upper arm, (4) forehead, (5) upper chest, and (6) stomach. (**b**) PPG signal magnitudes for NIR light on the aforementioned 6 locations for *n*=10 participants. HRV is derived from PPG, hence, PPG signal magnitudes are used in the analysis.

**Supplementary Fig 9.**
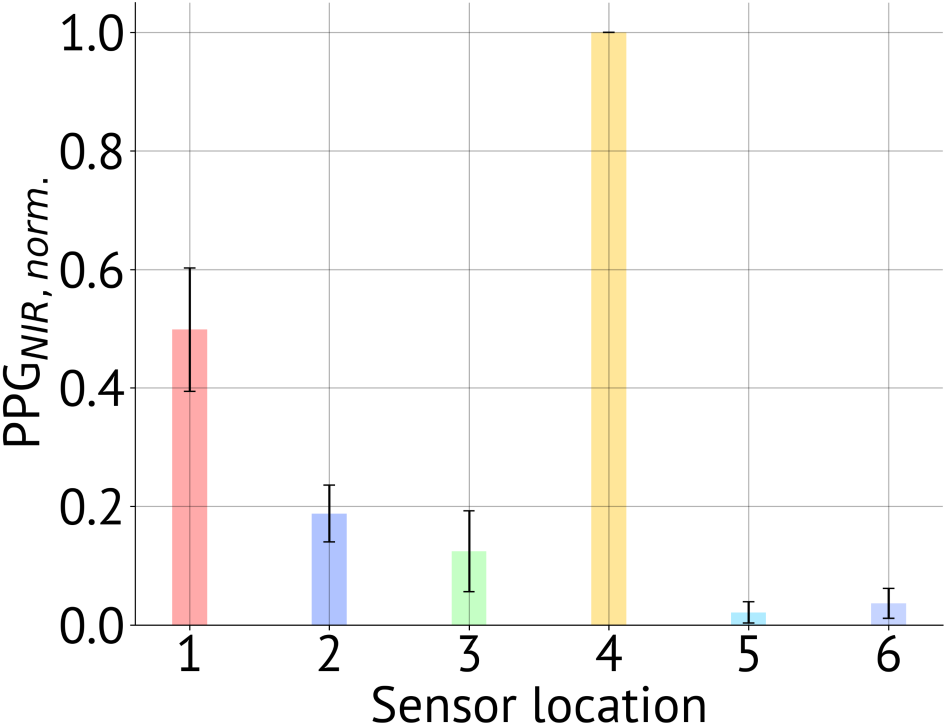
The reproducibility of HRV measurement. Since HRV is derived from PPG, PPG signal magnitudes are used in the analysis. Five consecutive NIR PPG measurements were collected from one participant while donning and doffing the sensor for each measurement on (1) wrist, (2) forearm, (3) upper arm, (4) forehead, (5) upper chest, and (6) stomach. NIR PPG signal was normalized for each set of measurements on the aforementioned six locations. The bar heights represent the average of the normalized value and the error bars represent the standard deviation of the normalized value.

**Supplementary Fig 10.**
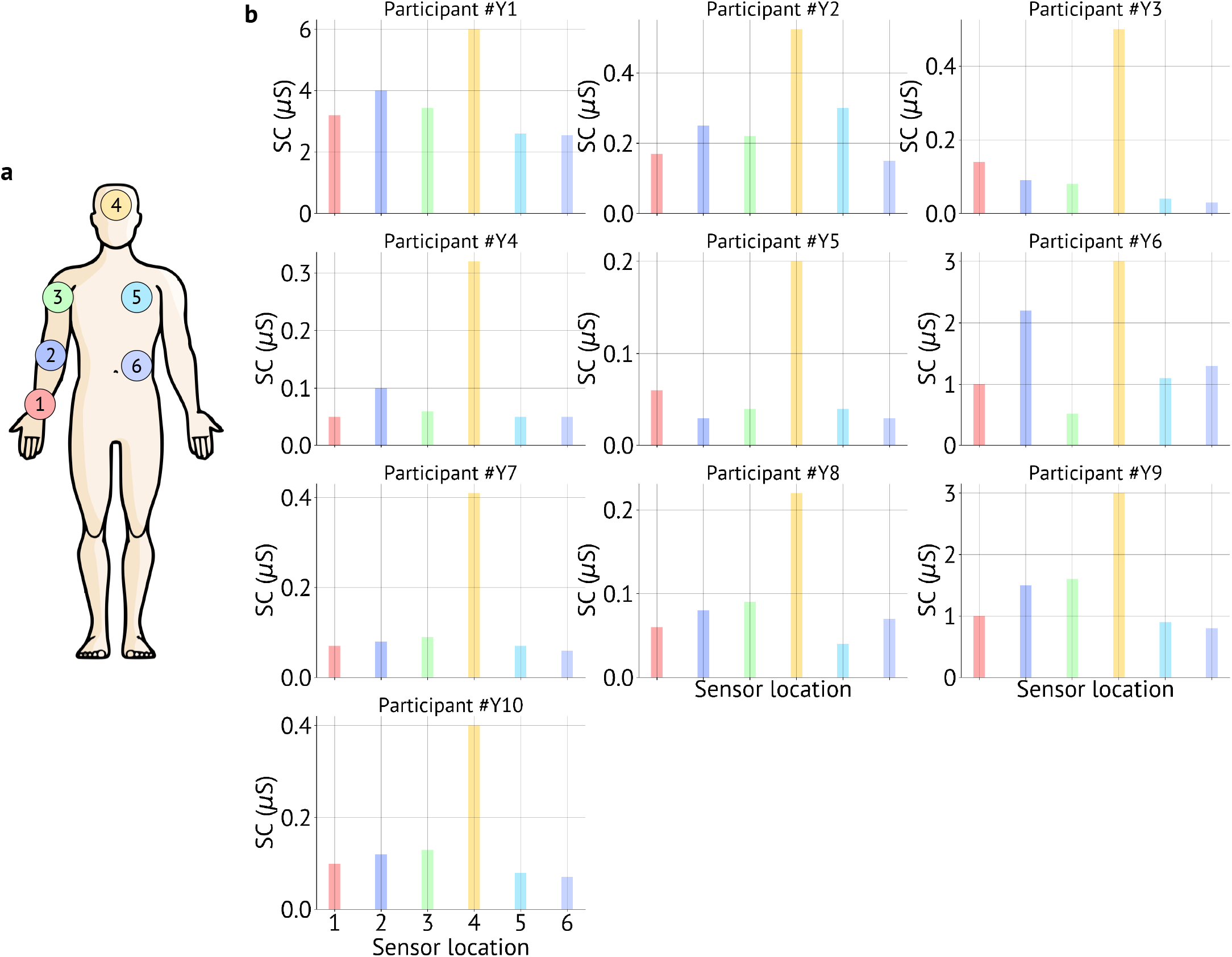
The complete SC dataset of *n*=10 participants. (**a**) Sensor placement locations − (1) wrist, (2) forearm, (3) upper arm, (4) forehead, (5) upper chest, and (6) stomach. (**b**) SC signal magnitudes on the aforementioned 6 locations for *n*=10 participants.

**Supplementary Fig 11.**
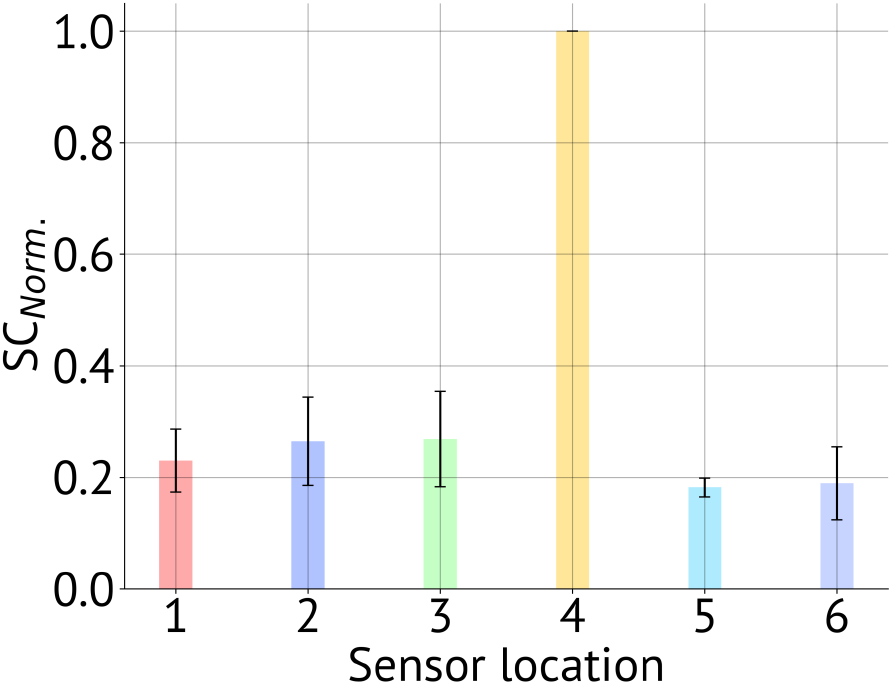
The reproducibility of SC measurement. Five consecutive SC measurements were collected from one participant while donning and doffing the sensor for each measurement on (1) wrist, (2) forearm, (3) upper arm, (4) forehead, (5) upper chest, and (6) stomach. SC signal was normalized for each set of measurements on the aforementioned six locations. The bar heights represent the average of the normalized value and the error bars represent the standard deviation of the normalized value.

**Supplementary Fig 12.**
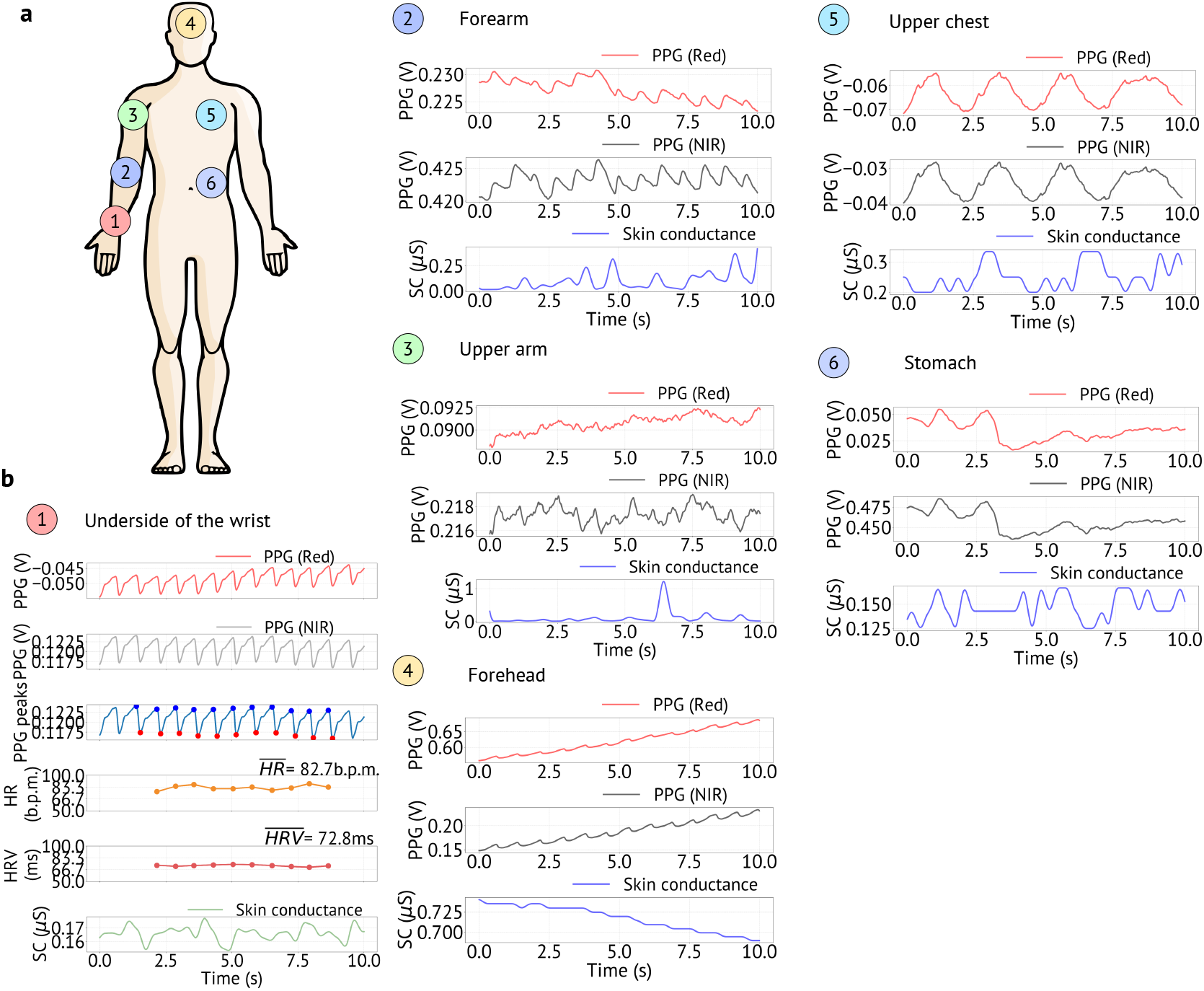
Impact of respiration on biosignals on the chest and stomach. (**a**) Heart rate variability (HRV) and skin conductance (SC) data collection locations − (1) wrist, (2) forearm, (3) upper arm, (4) forehead, (5) upper chest, and (6) stomach. (**b**) PPG from red and NIR channels, systolic and diastolic peaks from PPG, heart rate (HR), HRV calculated from PPG signal, and SC from the 6 highlighted locations shown in a. The PPG signal is clear on the wrist, forearm, upper arm, and forehead. The PPG signal gets highly attenuated on the upper chest and stomach. The modulation is obvious on the chest, and the respiration signal is clearly visible in b(5). Therefore, it is hard to get the PPG signal, hence, HRV cannot be calculated from the chest or the stomach.

### 2 Supplementary Note 2: Impact of respiration on biosignals on the chest and stomach

During the on-body biosignal data collection, we observed on the chest and the stomach, the PPG signal gets modulated with respiration. Since the PPG signal is a small pulsating signal on top of a big static signal, any motion gets coupled to both signals. A representative data where respiration severely affects the PPG signal is shown in Supplementary Fig. 12b(5)-(6). The modulation is obvious on the chest, and the respiration signal is clearly visible. In such a scenario, it is hard to get the PPG signal, hence, HRV cannot be calculated from the chest or the stomach.

**Supplementary Fig 13.**
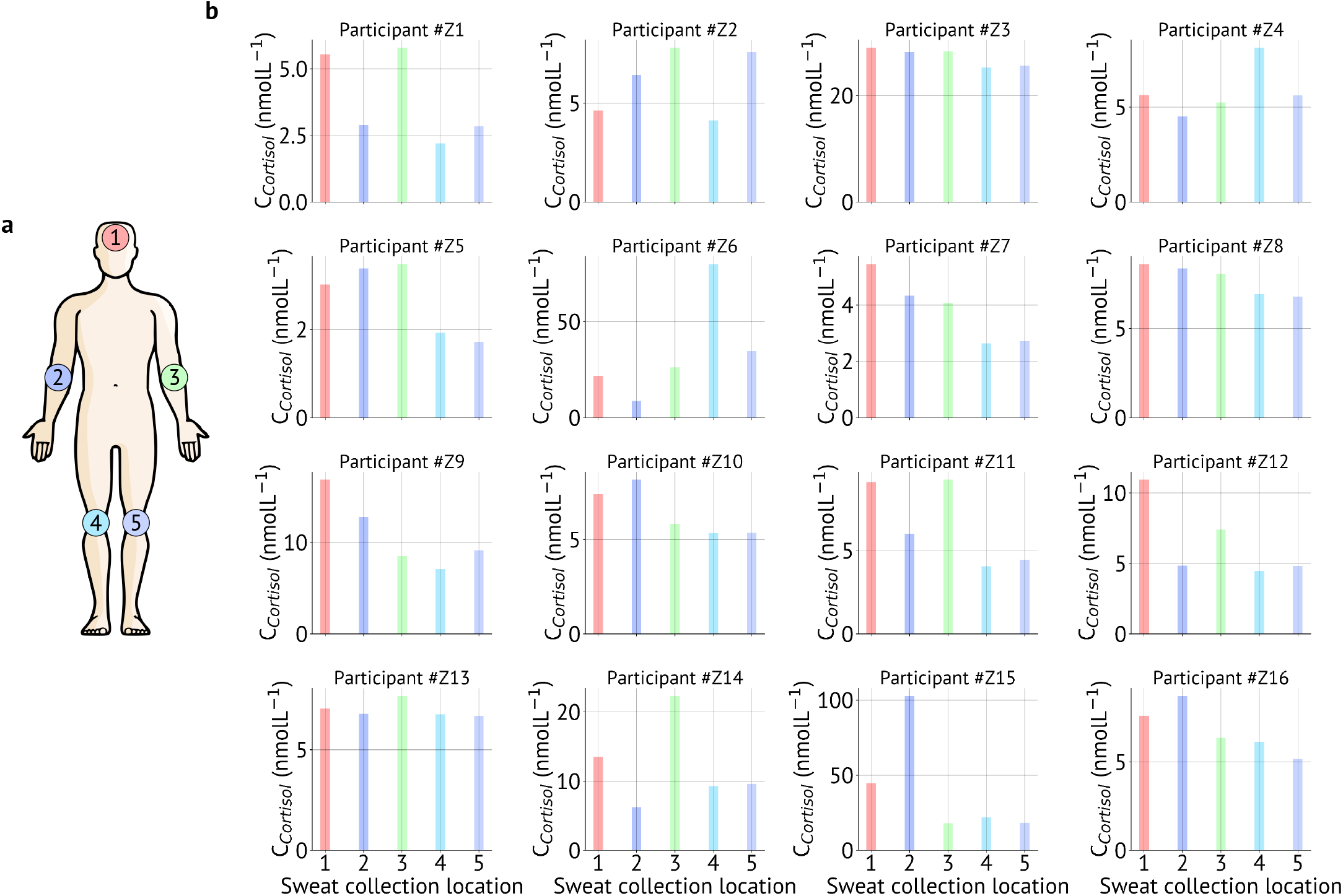
Sweat cortisol distribution on the body. (**a**) Sweat collection locations - (1) forehead, (2) right arm (cubital fossa), (3) left arm (cubital fossa), (4) back of the right knee (popliteal fossa), and (5) back of the left knee (popliteal fossa). (**b**) Sweat cortisol concentrations on the aforementioned 5 locations. The bar chart shows data collected from *n*=16 participants.

### 3 Supplementary Note 3: Individual impacts of SC and HRV on optimal wear location

In our analysis, the user preference data was collected from the design probe study, and the biosignals data was collected from the on-body sensing. Then, we combined SC, HRV, and cortisol body maps to create the biosignal body contour map using equal weights, *S*_*Biosignals*_ = *w*_1_ *× S*_*SC*_ + *w*_2_ *× S*_*HRV*_ + *w*_3_ *× S*_*Cortisol*_, where, *w*_1_ = *w*_2_ = *w*_3_ = 0:33. However, in this section, we investiage the effects of SC on *I*_*Wear*_using the equation: *I*_*Wear*_ = *w*_1_ *× S*_*Preference*_ + *w*_2_ *× S*_*SC*_ (Supplementary Figs. 14a-c). Similarly, we investiage the effects of HRV on *I*_*Wear*_ using the equation: *I*_*Wear*_ = *w*_1_ *× S*_*Preference*_ + *w*_2_ *× S*_*HRV*_ (Supplementary Figs. 14d-f). In both cases, *w*_1_ and *w*_2_ were assigned the combinations of (*w*_1_ = 0.75, *w*_2_ = 0.25), (*w*_1_ = 0.50, *w*_2_ = 0.50), and (*w*_1_ = 0.25, *w*_2_ = 0.75). It is important to mention that our HRV signal is derived from PPG signal. Therefore, we used PPG signal magnitude as a proxy for HRV signal.

For SC, with (*w*_1_ = 0.75, *w*_2_ = 0.25) and (*w*_1_ = 0.50, *w*_2_ = 0.50), *I*_*Wear*_ remains high on places that are normally covered with everyday clothing. For (*w*_1_ = 0.25, *w*_2_ = 0.75), *I*_*Wear*_ is high on the extermiteis of the body. For HRV, with (*w*_1_ = 0.75, *w*_2_ = 0.25) *I*_*Wear*_ remain high on places that are normally covered with everyday clothing. When, (*w*_1_ = 0.50, *w*_2_ = 0.50), *I*_*Wear*_ is high on the upper arm and the forearm areas. With (*w*_1_ = 0.25, *w*_2_ = 0.75), the body contour map almost resembles the Biosignals body map, and *I*_*Wear*_ is high on the extermiteis of the body. Note that, due to the high variablity of the HRV signal on the body, the effects of the HRV signal is more pronouced on the optimal placement location.

**Supplementary Fig 14.**
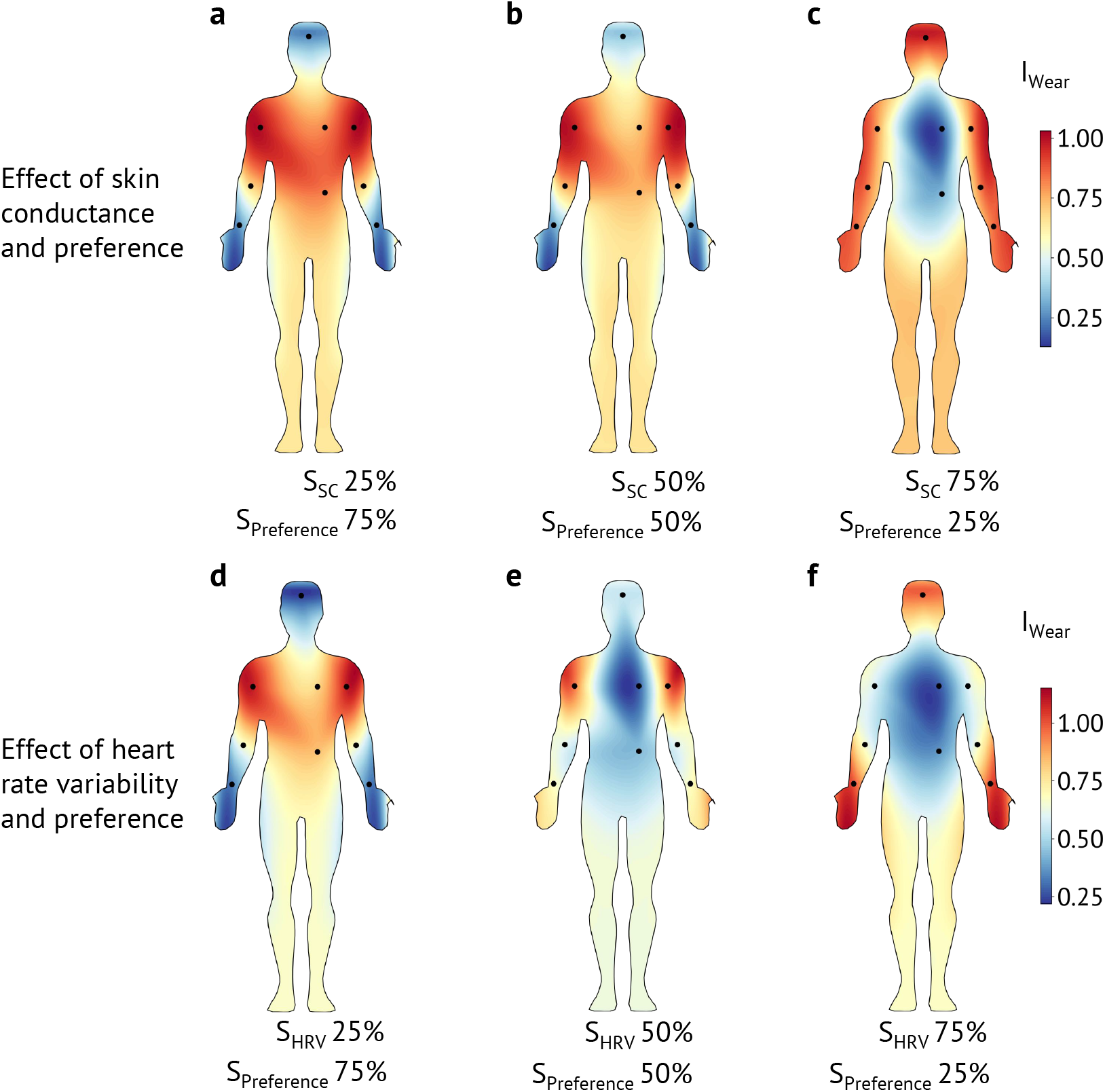
Individual impacts of SC and HRV on optimal wear location. (**a-c**) The implact of SC on *I*_*Wear*_ is investigated using the equation: *I*_*Wear*_ = *w*_1_ *× S*_*Preference*_ + *w*2 *× S*_*SC*_. Here, *w*_1_ and *w*_2_ were assigned the combinations of (*w*_1_ = 0.75, *w*_2_ = 0.25) in a, (*w*_1_ = 0.50, *w*_2_ = 0.50) in b, and (*w*_1_ = 0.25, *w*_2_ = 0.75) in c. (**d-f**) The implact of HRV on *I*_*Wear*_ is investigated using the equation: *I*_*Wear*_ = *w*_1_ *× S*_*Preference*_ + *w*2 *× S*_*HRV*_. Here, *w*_1_ and *w*_2_ were assigned the combinations of (*w*_1_ = 0.75, *w*_2_ = 0.25) in d, (*w*_1_ = 0.50, *w*_2_ = 0.50) in e, and (*w*_1_ = 0.25, *w*_2_ = 0.75) in f. Due to the high variablity of the HRV signal on the body, the effects of the HRV signal is more pronounced on the optimal placement location.

## References

[1] Almeida, D. M. Resilience and vulnerability to daily stressors assessed via diary methods. Current Directions in Psychological Science 14, 64–68 (2005).

[2] McEwen, B. S. Stress, adaptation, and disease: Allostasis and allostatic load. Annals of the New York academy of sciences 840, 33–44 (1998).

[3] Epel, E. S. et al. Accelerated telomere shortening in response to life stress. Proceedings of the National Academy of Sciences 101,17312–17315 (2004).

[4] Patel, V. et al. The lancet commission on global mental health and sustainable development. The Lancet 392, 1553–1598 (2018).

[5] Friedrich, M. J. Depression is the leading cause of disability around the world. Jama 317, 1517–1517 (2017).

[6] (WHO), W. H. O. et al. Suicide: One person dies every 40 seconds. Retrieved February 3, 2020 (2019).

[7] Kemp, A. H. et al. Impact of depression and antidepressant treatment on heart rate variability: a review and meta-analysis. Biological Psychiatry 67, 1067–1074 (2010).

[8] Chalmers, J. A., Quintana, D. S., Abbott, M. J., Kemp, A. H. et al. Anxiety disorders are associated with reduced heart rate variability: a meta-analysis. Frontiers in Psychiatry 5, 80 (2014).

[9] Alvares, G. A., Quintana, D. S., Hickie, I. B. & Guastella, A. J. Autonomic nervous system dysfunction in psychiatric disorders and the impact of psychotropic medications: a systematic review and meta-analysis. Journal of Psychiatry & Neuroscience (2016).

[10] Boucsein, W. Electrodermal activity (Springer Science & Business Media, 2012).

[11] Vahey, R. & Becerra, R. Galvanic skin response in mood disorders: A critical review (2015).

[12] Sarchiapone, M. et al. The association between electrodermal activity (eda), depression and suicidal behaviour: A systematic review and narrative synthesis. BMC Psychiatry 18, 22 (2018).

[13] Knorr, U., Vinberg, M., Kessing, L. V. & Wetterslev, J. Salivary cortisol in depressed patients versus control persons: a systematic review and meta-analysis. Psychoneuroendocrinology 35, 1275–1286 (2010).

[14] Stetler, C. & Miller, G. E. Depression and hypothalamic-pituitary adrenal activation: a quantitative summary of four decades of research. Psychosomatic Medicine 73, 114–126 (2011).

[15] Murri, M. B. et al. Hpa axis and aging in depression: systematic review and meta-analysis. Psychoneuroendocrinology 41, 46–62 638 (2014).

[16] Boggero, I. A., Hostinar, C. E., Haak, E. A., Murphy, M. L. & Segerstrom, S. C. Psychosocial functioning and the cortisol awakening response: Meta-analysis, p-curve analysis, and evaluation of the evidential value in existing studies. Biological Psychology 129, 207–230 (2017).

[17] Hogenelst, K., Soeter, M. & Kallen, V. Ambulatory measurement of cortisol: Where do we stand, and which way to follow? Sensing and Bio-Sensing Research 22, 100249 (2019).

[18] Valenza, G. et al. Wearable monitoring for mood recognition in bipolar disorder based on history-dependent long-term heart rate variability analysis. IEEE Journal of Biomedical and Health Infor650 matics 18, 1625–1635 (2013)

[19] Mohan, P. M., Nagarajan, V. & Das, S. R. Stress measurement from wearable photoplethysmographic sensor using heart rate variability data. In 2016 International Conference on Communication and Signal Processing (ICCSP), 1141–1144 (IEEE, 2016).

[20] Poh, M.-Z., Swenson, N. C. & Picard, R. W. A wearable sensor for unobtrusive, long-term assessment of electrodermal activity. IEEE Transactions on Biomedical Engineering 57, 1243–1252 (2010).

[21] Yoon, S., Sim, J. K. & Cho, Y.-H. A flexible and wearable human stress monitoring patch. Scientific Reports 6, 23468 (2016).

[22] Parlak, O., Keene, S. T., Marais, A., Curto, V. F. & Salleo, A. Molecularly selective nanoporous membrane-based wearable organic electrochemical device for noninvasive cortisol sensing. Science Ad663 vances 4, eaar2904 (2018).

[23] Torrente-Rodríguez, R. M. et al. Investigation of cortisol dynamics in human sweat using a graphene-based wireless mhealth system. Matter (2020).

[24] Niu, S. et al. A wireless body area sensor network based on stretchable passive tags. Nature Electronics 2, 361–368 (2019).

[25] Khan, Y., Ostfeld, A. E., Lochner, C. M., Pierre, A. & Arias, A. C.Monitoring of vital signs with flexible and wearable medical devices. Advanced Materials 28, 4373–4395 (2016).

[26] Chung, H. U. et al. Skin-interfaced biosensors for advanced wireless physiological monitoring in neonatal and pediatric intensive-care units. Nature Medicine 26, 418–429 (2020).

[27] Dunne, L., Profita, H. & Zeagler, C. Social aspects of wearability and interaction. In Wearable Sensors, 25–43 (Elsevier, 2014).

[28] Beauchaine, T. P. & Thayer, J. F. Heart rate variability as a trans-diagnostic biomarker of psychopathology. International Journal of Psychophysiology 98, 338–350 (2015).

[29] Steckl, A. J. & Ray, P. Stress biomarkers in biological fluids and their point-of-use detection. ACS Sensors 3, 2025–2044 (2018).

[30] Khan, Y. et al. A flexible organic reflectance oximeter array. Proceedings of the National Academy of Sciences 115, E11015– 683 E11024 (2018).

[31] Khan, Y. et al. Organic multi-channel optoelectronic sensors for wearable health monitoring. IEEE Access 7, 128114–128124 686 (2019).

[32] Bariya, M. et al. Glove-based sensors for multimodal monitoring of natural sweat. Science Advances 6, eabb8308 (2020).

[33] Taylor, N. A. & Machado-Moreira, C. A. Regional variations in transepidermal water loss, eccrine sweat gland density, sweat secretion rates and electrolyte composition in resting and exercising humans. Extreme Physiology & Medicine 2, 4 (2013).

[34] Zeagler, C. Where to wear it: functional, technical, and social considerations in on-body location for wearable technology 20 years of designing for wearability. In Proceedings of the 2017 ACM International Symposium on Wearable Computers, 150–157 (2017).

[35] Rüsch, N., Angermeyer, M. C. & Corrigan, P. W. Mental illness stigma: Concepts, consequences, and initiatives to reduce stigma. European Psychiatry 20, 529–539 (2005).

[36] Hinshaw, S. P. & Cicchetti, D. Stigma and mental disorder: Conceptions of illness, public attitudes, personal disclosure, and social policy. Development and Psychopathology 12, 555–598 (2000).

[37] Profita, H. P. et al. Don’t mind me touching my wrist: a case study of interacting with on-body technology in public. In Proceedings of the 2013 International Symposium on Wearable Computers, 89–96 706 (2013).

[38] Kelly, N. & Gilbert, S. The wear scale: Developing a measure of the social acceptability of a wearable device. In Proceedings of the 2016 CHI Conference Extended Abstracts on Human Factors in Computing Systems, 2864–2871 (2016).

[39] Kelly, N. The wear scale: Development of a measure of the social acceptability of a wearable device. Graduate Theses and Dissertations, Iowa State University (2016).

[40] Ingram IV, P. B.-, Clarke, E. & Lichtenberg, J. W. Confirmatory factor analysis of the perceived stress scale-4 in a community sample. Stress and Health 32, 173–176 (2016).

[41] Warttig, S. L., Forshaw, M. J., South, J. & White, A. K. New,normative, english-sample data for the short form perceived stress scale (pss-4). Journal of Health Psychology 18, 1617–1628 (2013).

## References

[1] Kelly, N. & Gilbert, S. The wear scale: Developing a measure of the social acceptability of a wearable device. In Proceedings of the 2016 CHI Conference Extended Abstracts on Human Factors in Computing Systems, 2864–2871 (2016).

[2] Kelly, N. The wear scale: Development of a measure of the social acceptability of a wearable device. Graduate Theses and Dissertations, Iowa State University (2016).

[3] Ingram IV, P. B., Clarke, E. & Lichtenberg, J. W. Confirmatory factor analysis of the perceived stress scale-4 in a community sample. Stress and Health 32, 173–176 (2016).

[4] Warttig, S. L., Forshaw, M. J., South, J. & White, A. K. New, normative, english-sample data for the short form perceived stress scale (pss-4). Journal of Health Psychology 18, 1617–1628 (2013).

